# (p)ppGpp/GTP and malonyl-CoA modulate *Staphylococcus aureus* adaptation to FASII antibiotics and provide a basis for synergistic bi-therapy

**DOI:** 10.1101/2020.03.26.007567

**Authors:** Amit Pathania, Jamila Anba-Mondoloni, Myriam Gominet, David Halpern, Julien Dairou, Laëtitia Dupont, Gilles Lamberet, Patrick Trieu-Cuot, Karine Gloux, Alexandra Gruss

## Abstract

Fatty acid biosynthesis (FASII) enzymes are considered valid targets for antimicrobial drug development against the human pathogen *Staphylococcus aureus*. However, incorporation of host fatty acids confers FASII antibiotic adaptation that compromises prospective treatments. *S. aureus* adapts to FASII inhibitors by a first non-replicative latency period followed by outgrowth. Here we used transcriptional fusions and direct metabolite measurements to investigate the factors that dictate the duration of latency prior to outgrowth. We show that stringent response induction leads to repression of FASII and phospholipid synthesis genes. (p)ppGpp induction inhibits synthesis of malonyl-CoA, a molecule that derepresses FapR, a key regulator of FASII and phospholipid synthesis. Anti-FASII treatment also triggers transient expression of (p)ppGpp-regulated genes during the anti-FASII latency phase, with concomitant repression of FapR regulon expression. These effects are reversed upon outgrowth. GTP depletion, a known consequence of stringent response, also occurs during FASII latency, and is proposed as the common signal linking these responses. We next show that anti-FASII treatment shifts malonyl-CoA distribution between its interactants FapR and FabD, towards FapR, increasing expression of phospholipid synthesis genes *plsX* and *plsC* during outgrowth. We conclude that components of the stringent response dictate malonyl-CoA availability in *S. aureus* FASII regulation, and contribute to latency prior to anti-FASII-adapted outgrowth. A combinatory approach, coupling a (p)ppGpp inducer and an anti-FASII, blocks *S. aureus* outgrowth, opening perspectives for bi-therapy treatment.

**Importance:** *Staphylococcus aureus* is a major human bacterial pathogen for which new inhibitors are urgently needed. Antibiotic development has centered on the fatty acid synthesis (FASII) pathway, which provides the building blocks for bacterial membrane phospholipids. However, *S. aureus* overcomes FASII inhibition and adapts to anti-FASII by using exogenous fatty acids that are abundant in host environments. This adaptation mechanism comprises a transient latency period, followed by bacterial outgrowth. Here we use metabolite sensors and promoter reporters to show that responses to stringent conditions and to FASII inhibition intersect: both involve GTP and malonyl-CoA. These two signaling molecules contribute to modulating the duration of latency prior to *S. aureus* adaptation outgrowth. We exploit these novel findings to propose a bi-therapy treatment against staphylococcal infections.

## Introduction

Bacterial infections that fail to respond to antibiotic treatments are on the rise, especially in the immunocompromised or weakened host, underlining the need for novel antimicrobial strategies (1). The fatty acid synthesis (FASII) enzymes were considered as fail-safe targets for eliminating numerous gram-positive pathogens. Anti-FASII has been a front-line drug against *Mycobacterium tuberculosis*, which synthesize very long chain fatty acids that cannot be compensated by the environment (2). However, Firmicute pathogens, including *Staphylococcus aureus* and numerous streptococcaceae, bypass FASII inhibition and satisfy their fatty acid requirements by using host-supplied fatty acids (3–5). FASII inhibitors like triclosan (Tric), MUT056399, fasamycins A and B, amycomicin and a pipeline FASII antibiotic AFN-1252 (6–11), would thus have limited use as stand-alone treatments of infections by numerous gram-positive pathogens (3–5).

Our recent studies show that *S. aureus* can adapt to FASII inhibitors by two mechanisms, depending on growth conditions. One involves mutations in a FASII initiation gene, usually *fabD*. Lower activity of the FabD mutant would increase availability of its substrates, one of which is acyl carrier protein (ACP), for incorporation of exogenous fatty acids (eFA) *via* the phosphate acyltransferase PlsX (**Fig. S1A**) (4, 12). The second mode of adaptation occurs without FASII mutations and predominates in serum-supplemented medium. In this case, full adaptation and eFA incorporation in actively growing cells is achieved after a latency phase, whose duration (6-12 h) depends on the strain and pre-growth in serum-containing medium. Adaptation is associated with greater intracellular retention of eFA and ACP, which both contribute to eFA incorporation in membrane phospholipids to compensate FASII inhibition (**Fig. S1B**(5)).

The factors regulating *S. aureus* transition from latency to outgrowth upon anti-FASII treatment remain unknown. We hypothesized that initial fatty acid starvation in response to anti-FASII might comprise the signal that delays eFA incorporation in phospholipids and outgrowth. The *S. aureus* FapR repressor reportedly regulates most FASII genes (except *acc*, encoding acetyl-CoA carboxylase, and FabZ, β-hydroxyacyl-ACP dehydratase), and phospholipid synthesis genes *plsX* and *plsC* (13, 14). Interestingly, malonyl-CoA has a dual function: it is the first dedicated FASII substrate used by FabD (malonyl-CoA transacylase), and it also controls FapR by a feed-forward mechanism (14). FabD uses malonyl-CoA and ACP to synthesize malonyl-ACP (15). Malonyl-CoA binding to FapR reverses FapR repression, leading to up-regulation of the FASII and phospholipid synthesis genes (14). Thus, malonyl-CoA is important in both enzymatic and regulatory activities of FASII. In *Escherichia coli*, expression of the malonyl-CoA synthesis enzyme ACC is regulated by (p)ppGpp, which accumulates in slow growing, nutrient-deficient conditions (16, 17); (p)ppGpp also reportedly regulates other FASII and phospholipid synthesis genes (18, 19). In *B. subtilis*, studies of (p)ppGpp null mutants gave evidence for the need to activate stringent response in order to survive fatty acid starvation; these studies implicated increased GTP in mortality of (p)ppGpp null mutant strains (20). Fatty acid starvation is also associated with cell size *via* regulation of FASII, although underlying mechanisms remain to be elucidated (21). To our knowledge, no evidence exists for stringent response-mediated FASII regulation in *S. aureus*.

Here, we first show that stringent response induction exerts control over fatty acid and phospholipid synthesis in *S. aureus* by modulating FapR repressor activity. FASII antibiotic treatment, like stringent response, leads to GTP depletion, which is the likely common metabolite linking these two responses. The chain of events revealed here indicate that (p)ppGpp/GTP and malonyl-CoA contribute to adjusting the timing of FASII-antibiotic-induced latency transition to outgrowth. Based on our findings, we suggest a bi-therapy approach that combines FASII inhibitors and a (p)ppGpp inducer to prevent *S. aureus* adaptation.

## Results

### (p)ppGpp negatively regulates malonyl-CoA levels in *S. aureus*

We investigated the potential roles of (p)ppGpp and malonyl-CoA in *S. aureus* response to FASII inhibition. Three *S. aureus* strains were used in this study, Newman, USA300, and HG1-R (**Table S1**), which all adapt to anti-FASII with similar kinetics ((5), this study). Previous studies reported difficulties in (p)ppGpp measurements in *B. subtilis* and *S. aureus* (20, 22). Our initial attempts at measuring (p)ppGpp by HPLC and fluorescent dye PyDPA (23) failed to give reliable results (data not shown). We therefore constructed transcriptional fusions to detect conditions when (p)ppGpp-induced genes are activated *in vivo* (**Table S2**). The reporter fusion activities responded to mupirocin, which inhibits isoleucyl-tRNA synthetase and triggers (p)ppGpp synthesis (24) (**Fig. 1A** and data not shown). P_*ilvD*_-*lacZ* (NWMN_1960) and P_*oppB*_-*lacZ* (NWMN_0856) were up-regulated, and P_*cshA*_-*lacZ* (NWMN_1985) was down-regulated by mupirocin. Nutrient starvation during stationary phase induces stringent response in *E. coli* (16). In *S. aureus*, β-galactosidase (β-gal) activity of the P_*ilvD*_-*lacZ* and P_*oppB*_-*lacZ* sensors were 1.2- and 7-fold higher in stationary phase compared to exponential phase cells, while P_*cshA*_-*lacZ* activity was ^~^2-fold lower (**Fig. 1B** and data not shown), further validating the *in vivo* (p)ppGpp sensors.

**Fig. 1.**
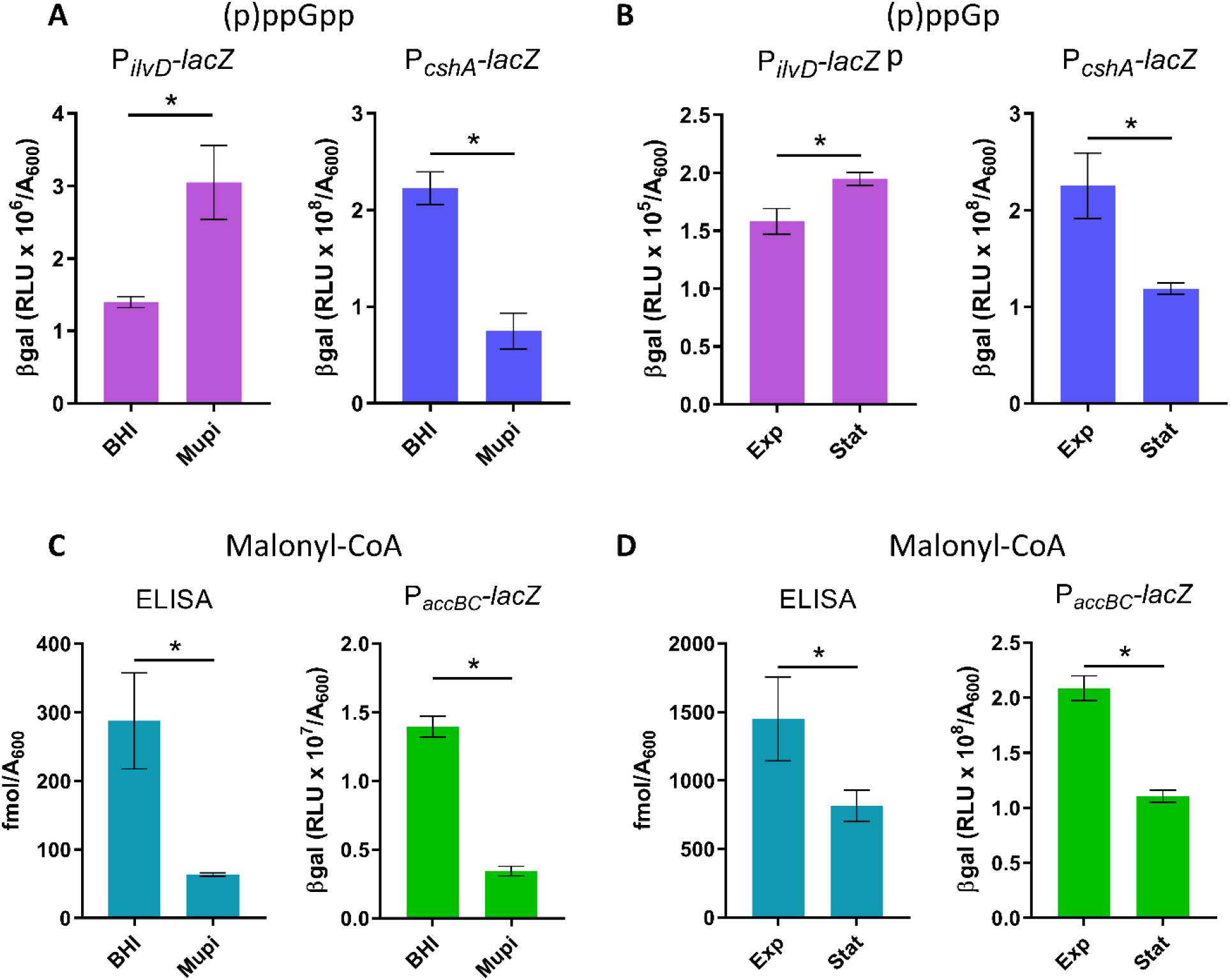
Mupirocin and stationary phase conditions stimulate (p)ppGpp sensor responses and inhibit malonyl-CoA production and *accBC* activity. *S. aureus* Newman strains contain reporter systems as indicated. Strains were grown in BHI and BHI containing 0.1 μg/ml mupirocin (Mupi) for 1h (**A**,**C**), or in SerFA (*A*_600_ ~4.0, Exp) and stationary phase (*A*_600_ ~10.0, Stat) (**B** and **D**). P_*ilvD*_-*lacZ* and P_*cshA*_-*lacZ* expression were evaluated by β-gal assays. Total Malonyl-CoA levels were determined by immunoassay (ELISA), and deduced from P_*accBC*_-*lacZ* expression. *ilvD* and *cshA* genes are respectively up-regulated and down-regulated by stringent response induction. Data presented are mean ± standard deviation from triplicate independent experiments. *, p ≤ 0.05 using Mann Whitney.

The stringent response sensors would expectedly not respond to mupirocin in a (p)ppGpp null strain. We compared sensor responses in HG1-R and HG1R-ppGpp0 strains. These strains derive from HG001 (25) and a (p)ppGpp-null strain (kindly provided by C. Wolz; (26)). They were first repaired for a defect in *fakB1*, which is common to 8325 derivatives (like HG001) and a minority of *S. aureus* isolates (5). FakB1, a fatty acid kinase subunit, facilitates assimilation of mainly saturated fatty acids (27). Its absence in 8325 derivatives can explain previous reports of *S. aureus* sensitivity to anti-FASII (11, 28), although the majority of *S. aureus* strains adapt to these antibiotics (4, 5). The *fakB1*-repaired HG001 and HG001 (p)ppGpp0 strains are referred to as respectively HG1-R and ppGpp0. Responses of the P_*ilvD*_-*lacZ*, and P_*cshA*_-*lacZ* reporter fusions were compared in HG1-R and ppGpp0 strains by plate tests (**Figure S2; see Materials and Methods)**. If the response to mupirocin occurs *via* its stimulation of (p)ppGpp, then neither induction of *ilvD* nor suppression of *cshA* would occur in the ppGpp0 background. Indeed, P_*ilvD*_-*lacZ* (**Fig. S2A**) and P_*cshA*_-*lacZ* (**Fig. S2B**) responded to mupirocin as expected in the parental strain, whereas no such responses were observed in the ppGpp0 background. These results also indicate that stringent response controls these sensors in *S. aureus*.

We then asked whether (p)ppGpp blocks malonyl-CoA synthesis in *S. aureus*, as reported in *E. coli* (29), despite major regulatory differences between these bacteria. Total malonyl-CoA was measured in cells treated or not with mupirocin by ELISA immunoassay. We also used *in vivo* promoter fusion P_*accBC*_-*lacZ* to measure expression of *accBC* (NWMN_1432 and NWMN_1431), which encode subunits of acetyl-CoA carboxylase (ACC) required for malonyl-CoA synthesis (**Table S2**). Stringent response induction by mupirocin led to decreases in malonyl-CoA pools (^~^6-fold) and in P_*accBC*_-*lacZ* β-gal activity (^~^4-fold) (**Fig. 1C**). Similarly, stationary phase cells showed ^~^2-fold lower malonyl-CoA production and P_*accBC*_-*lacZ* β-gal activity compared to exponential phase cells (**Fig. 1D**). Finally, the P_*accBC*_-*lacZ* reporter was inhibited by mupirocin in HG1-R, but not in the ppGpp0 strain (**Fig. S2C**). These results show that in *S. aureus*, stringent response induction leads to repression of malonyl-CoA synthesis (13).

### FASII-antibiotic-induced latency transiently alters expression of (p)ppGpp-regulated sensors

We recently showed that host fatty acids can compensate FASII-antibiotic inhibition of *S. aureus* to promote growth. In low membrane stress conditions as in serum, adaptation involves a transient latency phase without detection of FASII mutations (**Fig. 2A**). Anti-FASII-adapted *S. aureus* display fatty acid profiles that are fully exogenous (**Fig. S1B**, (5)). As anti-FASII treatment may provoke fatty acid deprivation before eFAs are incorporated, we asked whether the latency preceding FASII bypass corresponds to stringent response induction. Using the stringent response sensors, a ^~^3.9-fold increase in P_*ilvD*_-*lacZ* and ^~^7-fold decrease of P_*cshA*_-*lacZ* β-gal activities were observed during the latency phase preceding outgrowth (**Fig. 2B**), indicating that a factor related to stringent response is induced in response to anti-FASII treatment. P_*ilvD*_-*lacZ* activity returned to normal levels once bacteria are in the outgrowth phase. P_*cshA*_-*lacZ* β-gal activity was only partially restored during outgrowth, as levels increased by only 2-fold compared to latency. The reason for lower *cshA* expression is unknown, but it is likely that its expression is subject to other layers of regulation.

**Fig. 2.**
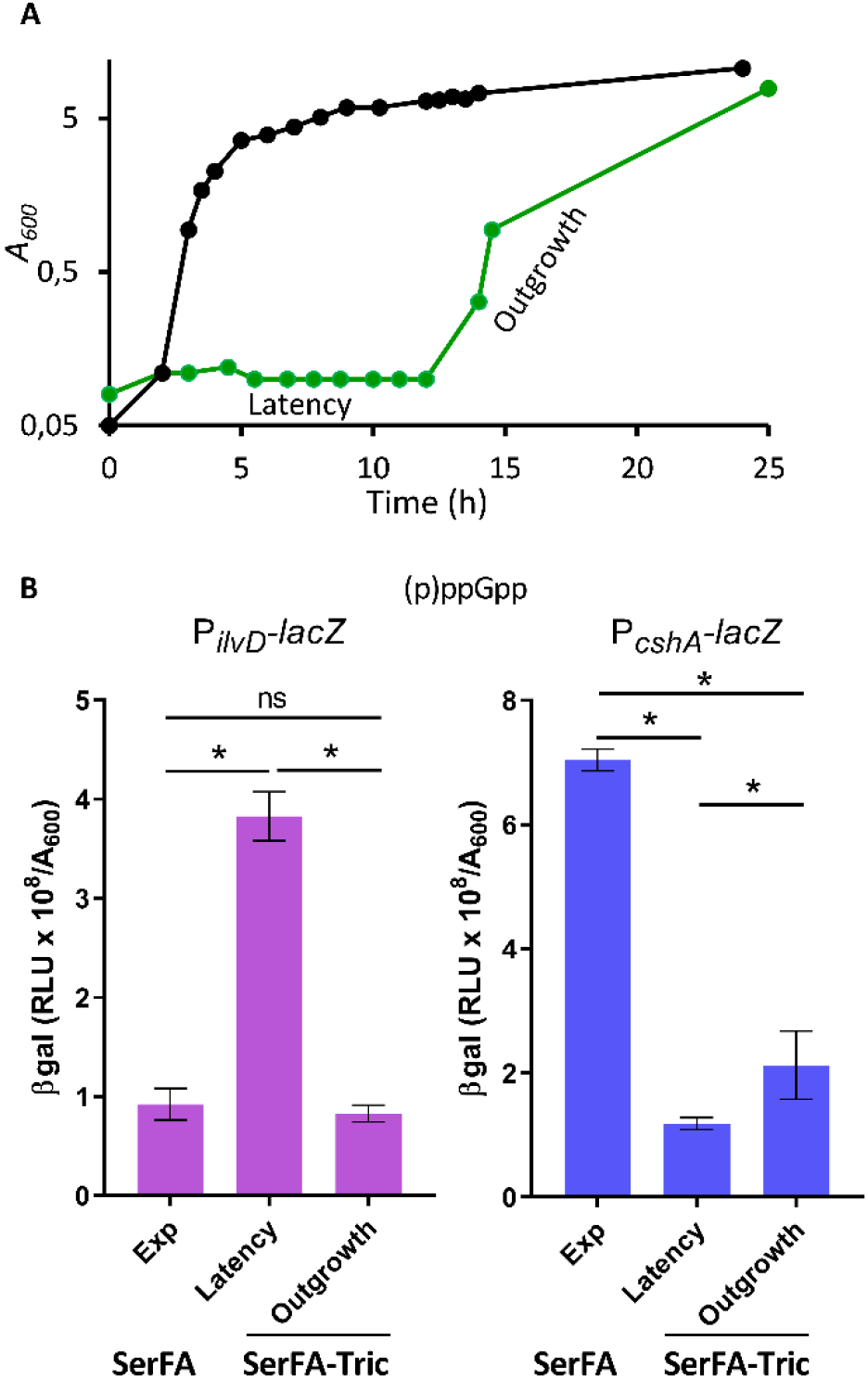
Anti-FASII treatment leads to transient responses of (p)ppGpp sensors. **A.** Growth kinetics of *S. aureus* Newman in SerFA (black line) and in SerFA-Tric (green). In SerFA-Tric, a 10-12 h latency period (Latency) precedes exponential outgrowth (Outgrowth). The growth curves are representative of three independent experiments. BHI and SerFA growth curves are essentially identical; no growth is observed in BHI medium containing triclosan without fatty acids (n=3, not shown). **B.** Stringent response status according to growth condition in Newman carrying P_*ilvD*_-*lacZ* reporter (left panel) and P_*cshA*_-*lacZ* (right panel). β-gal activities of samples in SerFA (exponential growth; Exp, *A*_600_ = ^~^1.5) and SerFA-Tric (Latency, *A*_600_ = ^~^0.3) are measured after 3 h growth. β-gal activities were measured on 17 h samples in SerFA-Tric (exponential growth; Outgrowth, *A*_600_ = ^~^1.5). Data presented are mean ± standard deviation from triplicate independent experiments. *, p ≤ 0.05, ns, not significant, using Mann Whitney.

### FASII antibiotic treatment down-regulates *accBC* and lowers malonyl-CoA pools

Malonyl-CoA, the ACC product, binds FapR and antagonizes repression, and is also a FabD substrate (**Fig. 3A**). We assessed malonyl-CoA production in non-selective (SerFA) and anti-FASII-treated (SerFA-Tric) latency and outgrowth Newman cultures. Pools of malonyl-CoA were measured by ELISA and by P_*accBC*_-*lacZ* expression. Both measurements indicated that malonyl-CoA levels were comparable in SerFA and SerFA-Tric-adapted outgrowth cultures, and were markedly lower during SerFA-Tric latency (**Fig. 3B**). Taken together these results show that stringent response induction and anti-FASII-induced latency lead to *accBC* inhibition, suggesting that a common element links these responses.

**Fig. 3.**
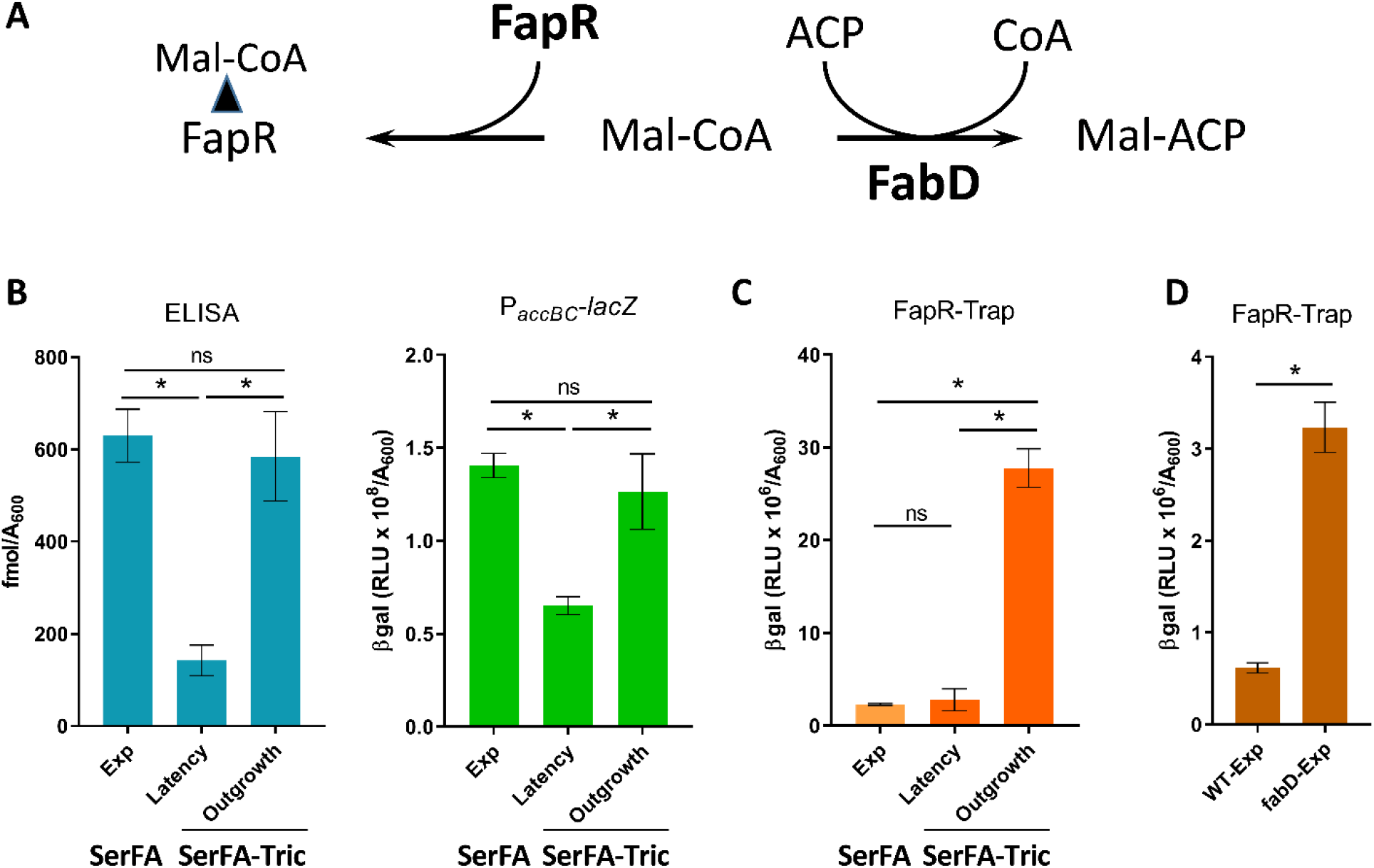
Anti-FASII treatment leads to transient *acc* repression and shifts in malonyl-CoA distribution. **A.** The diagram illustrates two known malonyl-CoA interactants, FapR and FabD. FapR is a repressor of FASII and phospholipid synthesis. Malonyl-CoA binds FapR (left), which reverses FapR repressor activity. FabD uses malonyl-CoA to produce malonyl-ACP, the FASII precursor (right). **B.** Assessment of malonyl-CoA production by ELISA and P_*accBC*_-*lacZ*. Left panel: Sandwich ELISA assay was used to measure total malonyl-CoA (Materials and Methods). Right panel: P_*accBC*_-*lacZ* reporter activity. **C.** Assessment of FapR-bound malonyl-CoA. FASII-antibiotic-adapted *S. aureus* display altered malonyl-CoA distribution (**see A**). Newman strain and derivatives were grown in non-selective (SerFA) and anti-FASII (SerFA-Tric) media. β-gal activities were measured in SerFA at *A*_600_ = ^~^2.0 (exponential growth: Exp), and in SerFA-Tric during latency after 3 h growth *A*_600_ = ^~^0.3 (Latency) and upon adaptive exponential outgrowth *A*_600_ = ^~^2 (Outgrowth). **D.** Assessment of FapR-bound malonyl-CoA in a *fabD* mutant using FapR-Trap. β-gal assays of Newman and *fabD* mutant (CondT^R^-17, a point mutant (4) carrying FapR-Trap were performed in SerFA after 3 h growth with *A*_600_ = ^~^1.0 (Exp). Measurements in **B** to **D** are mean ± standard deviation from triplicate independent experiments. *, p ≤ 0.05, ns, not significant, using Mann Whitney.

We also assessed pools of malonyl-CoA using a FapR activity sensor called FapR-Trap (**Fig. S3A, Table S2**). FapR-Trap responded as expected: expression was increased in the absence of repressor (Δ*fapR*), but decreased in stationary phase wild type cells when malonyl-CoA levels are low (**Fig. S3B**). Interestingly, and in sharp contrast to the above results, malonyl-CoA estimations by FapR-Trap were around 10-fold higher during SerFA-Tric outgrowth, compared to non-selective SerFA cultures (**Fig. 3C**; compare with **3B**). These differences (summarized in **Table S3**), particularly visible during adaptation outgrowth, indicate that malonyl-CoA distribution in anti-FASII-treated *S. aureus* favors FapR binding over FabD. They suggest that malonyl-CoA pools and their distribution between FapR and FabD may be central determinants in *S. aureus* adaptation to FASII antibiotics.

Reduced FabD competition for malonyl-CoA would increase its availability for FapR (**Fig. 3A**). We showed previously that *fabD* mutants may emerge upon FASII-antibiotic selection, but not in serum-supplemented medium as used here (4, 5). Indeed, a *fabD* mutant displayed 5-fold greater FapR-Trap expression than the parental strain in non-selective SerFA (Fig. 3D). However, we ruled out the presence of *fabD* mutations in our conditions by DNA sequencing of 5 independent anti-FASII-adapted cultures (available upon request). These findings could suggest that FabD is intact, but disabled for its interactions with malonyl-CoA during *S. aureus* growth in the presence of anti-FASII. This possibility is currently under study in our laboratory.

### GTP depletion is the feature common to stringent response and FASII-antibiotic-induced latency

We asked whether stringent response effector (p)ppGpp was directly responsible for the observed phenotypes during anti-FASII treatment, using a *S. aureus* WT (HG1-R) and the (p)ppGpp0 isogenic strain. AFN-1252 was used as anti-FASII in this strain background due to higher resistance of HG001 derivatives to triclosan. The HG1-R and (p)ppGpp0 strains grew similarly in the presence of anti-FASII, suggesting that the absence of (p)ppGpp did not accelerate anti-FASII adaptation (data not shown). We then compared expression of P_*ilvD*_-*lacZ* and P_*accBC*_-*lacZ* sensors in the WT *versus* (p)ppGpp0 backgrounds upon anti-FASII treatment (**Fig 4A**). Both sensors behaved as above (**Figs 2 and 3**) in the WT strain. However, these sensors displayed the same responses to anti-FASII treatment in the two strains. Thus, while (p)ppGpp induction inhibits *acc* and thus lowers malonyl-CoA pools, it is not required for these phenotypes in anti-FASII-treated *S. aureus*.

**Fig. 4.**
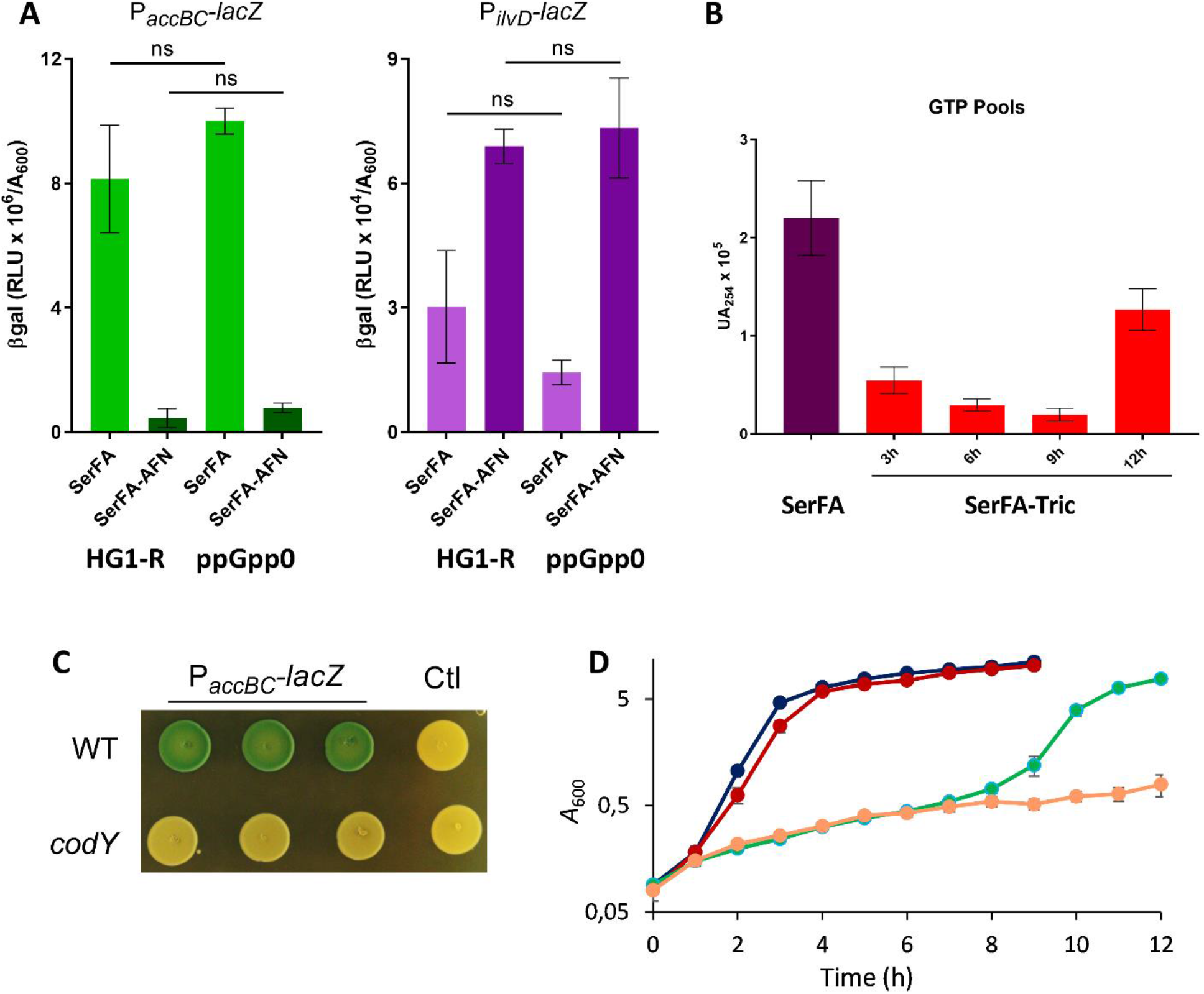
GTP is depleted during anti-FASII latency phase. **A.** P_*accBC*_-*lacZ* (left) and P_*ilvD*_-*lacZ* (right) sensor responses to anti-FASII latency were compared in *S. aureus* HG1-R and the (p)ppGpp null isogenic strain as indicated. β-gal activities were measured after 3h incubation in SerFA and in medium containing the anti-FASII AFN-1252. Data presented are mean ± standard deviation from three biological replicates. P-values were determined pairwise by Mann Whitney; ns, not significant. **B.** Newman strain GTP levels were assessed at different growth times during anti-FASII latency (3h, 6h, and 9h) and outgrowth (12h). Data presented are mean ± standard deviation from duplicate independent experiments. **C.** P _*accBC*_-*lacZ* expression is lower in a *codY* mutant. USA300 and *codY* derivative contain plasmids expressing P_*accBC*_-*lacZ* or the control plasmid (pTCV-lac; Ctl). Exponential phase cultures issued from three independent colonies were adjusted to A_600_ = 0.1 and 5μl drops were plated on BHI plates containing erythromycin (5 μg/ml) and X-gal. Photographs were taken after 20 h at 37°C and 24 h at 4°C. **D.** Growth of *S. aureus* USA300 and a confirmed *codY* mutant of the Nebraska mutant collection was compared in non-selective (SerFA) and SerFA-Tric conditions in four independent replicates. WT in SerFA, black; *codY* in SerFA, red; WT in SerFA-Tric, green; *codY* in SerFA-Tric, orange. Mean and standard deviation are shown for each time point.

(p)ppGpp is known to be intimately linked to GTP: (p)ppGpp inhibits GTP synthesis (30, 31). Lowering GTP rescues *B. subtilis* from ppGpp0 toxicity during lipid starvation(20). We used HPLC to measure GTP levels during anti-FASII adaptation of *S. aureus* Newman. GTP levels decreased by 4-fold at 3h post-anti-FASII treatment (**Fig. 4B**). Consistently, amounts of two GTP synthesis enzymes were decreased during anti-FASII latency of *S. aureus* USA300, as seen by proteomics (5); HprT (2.35-fold lower [n=4]; p=0.014) and GuaA (1.5-fold lower [n=4]; p=0.029). These results identify GTP as the metabolite and potential effector common to both stringent response and anti-FASII-induced latency.

GTP is also a cofactor of the pleiotropic regulator CodY (31). We asked whether CodY is implicated in *accBC* regulation. P_*accBC*_-*lacZ* expression was visibly lower in a *codY* insertional mutant compared to expression in the parental WT (USA300) **(Fig. 4C).** In addition, the anti-FASII latency period was strikingly longer in a *codY* mutant than in the WT strain (**Fig. 4D**). This delay is consistent with a role of GTP depletion in delaying anti-FASII latency *via* CodY. These results lead us to propose that in *S. aureus*, the stringent response pathway intersects the initial latency response to FASII inhibitors by the common depletion of GTP, likely *via* the CodY regulon.

### Phospholipid synthesis genes *plsX* and *plsC* are differently controlled by FapR

The above results show that malonyl-CoA pools are restored during *S. aureus* adaptation to FASII antibiotics, and preferentially bind FapR, which alleviates FapR repression (Fig. 3). The *S. aureus* FapR regulon reportedly includes *plsX* (NWMN_1139, part of the *fapR* operon) and *plsC* (NWMN_1620); however, the *S. aureus* FapR binding site in the *plsC* promoter region is highly degenerate ((13); see Fig. 5A), and no proof was given for this interaction. We used promoter reporter fusions P_*fapR plsX*_-*lacZ* and P_*plsC*_-*lacZ* (Table S2) to compare expression in a wild type strain (HG1-R) and its Δ*fapR* derivative. Expression of both reporters was upregulated (each 1.6-fold) in the Δ*fapR* strain (Fig. 5B). To determine whether regulation involved direct FapR binding, we performed DNase I footprinting using the *plsX* and *plsC* promoters as binding substrates for purified FapR (Fig. 5C). FapR bound efficiently to the *plsX* promoter region. In contrast, FapR did not bind the *plsC* upstream region containing the putative binding site. Taken together, these results indicate that in *S. aureus*, FapR regulates expression of both *plsX* and *plsC*, but that its effect on *plsC* is either indirect or may require other *S. aureus* factors.

**Fig. 5.**
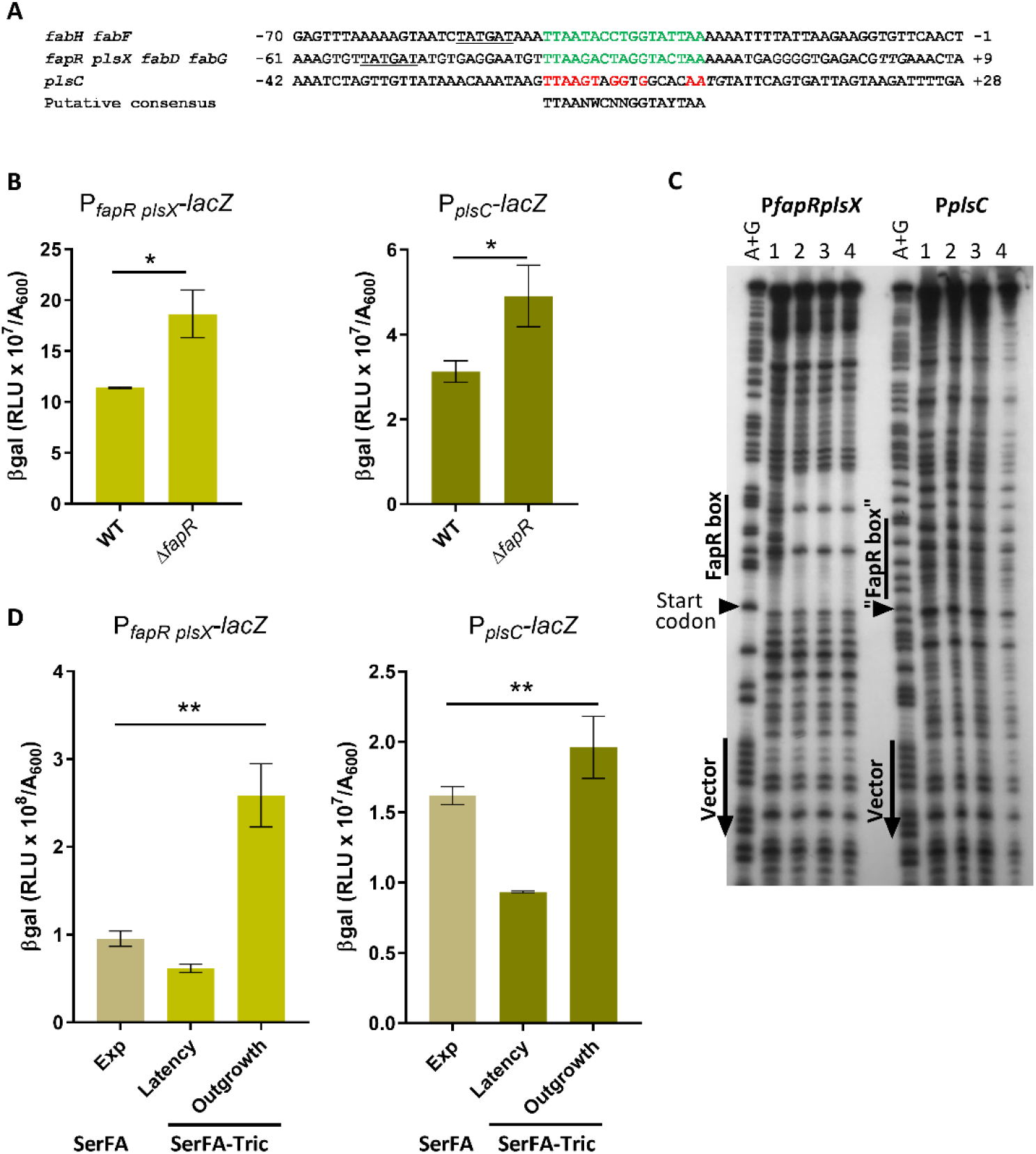
FapR binds the *plsX* but not the *plsC* promoter region, but affects expression of both genes. **A.** Sequence alignment of FapR binding sites in *S. aureus* as published (based on (13)). Positions correspond to the predicted start codon of the first Orf in each operon. Confirmed 17 bp FapR binding sequences are in green. The presumed conserved FapR binding nucleotides upstream of the *plsC* start site are in red. In the consensus sequence below the alignment: N, any nucleotide; W, A or T; Y, T or C. Putative −10 RNA polymerase binding sites are underlined (13). The PlsC ATG and FapR TTG start sites are in italics. **B**. Expression of P_*fapR plsX*_-*lacZ* and P_*plsC*_-*lacZ* fusions is upregulated in the Δ*fapR* strain. Strains were grown to *A*_600_ = ^~^1.5 in SerFA medium prior to β-gal determinations. Data presented are mean ± standard deviation from three biological replicates. *, p ≤ 0.05, ns, not significant, using Mann Whitney. **C.** DNase I footprinting assay of the *PfapRplsX* and P*plsC* promoter regions with FapR. Radiolabeled PCR fragments corresponding to P*plsX* and P*plsC* were used as DNA targets. Various amounts of FapR (lane 1: 0; lane 2: 1.2 nmol; lane 3: 6 nmol; and lane 4: 12 nmol) were incubated with 0.2 pmol DNA before DNase 1 digestion. Lane A+G: Maxam and Gilbert reaction of the labeled strand. Positions of vector and “FapR box” sequences are indicated by vertical lines, PlsX and PlsC start codons are indicated by black arrows. **D.**Expression of P_*fapR plsX*_-*lacZ* and P_*plsC*_-*lacZ* fusions in *S. aureus* Newman during growth in non-selective (SerFA) and anti-FASII (SerFA-Tric) media. β-gal are shown for samples processed in SerFA at OD_600_ = ^~^2.0 (exponential phase, Exp), and in SerFA-Tric after 3h in latency (OD_600_ = ^~^0.3) and upon adaptation outgrowth at 17 h (OD_600_ = ^~^2.0; Outgrowth). Mean and standard deviation are shown for three independent experiments. P-values were determined by Kruskal Wallis; p ≤ 0.005 is indicated by **; ns, not significant.

### Mupirocin and anti-FASII treatment lead to reduced expression of *S. aureus* phospholipid synthesis genes *plsX* and *plsC*

Repression of *accBC* FASII by mupirocin would expectedly impact all FapR-regulated genes, including those involved in phospholipid synthesis (**Fig. S1A**). To test this, we followed P_*fapRplsX*_-*lacZ* and P_*plsC*_-*lacZ* transcriptional fusion expression in the presence of mupirocin (0.1 μg/ml), using P_*ilvD*_-*lacZ* and P_*accBC*_-*lacZ* sensors as references (**Table S4**). Expression of *plsX* and *plsC* sensor fusions were 4- and 3-fold respectively lower in mupirocin than in non-treated samples.

Responses of P_*fapRplsX*_-*lacZ* and P_*plsC*_-*lacZ* during anti-FASII-induced latency and outgrowth were then measured. β-gal expression from both sensors gradually decreased during latency, followed by abrupt (respectively 4- and 2-fold) increases upon restart of active growth of anti-FASII -adapted cells (**Fig. 5D**, and data not shown). Expression of P_*fapRplsX*_-*lacZ* reached higher (^~^3-fold) levels in anti-FASII-adapted outgrowth than in non-selective growth. Anti-FASII treatment thus decreases expression of phospholipid synthesis genes during latency, which recovers upon adaptation.

### Mupirocin treatment lowers fatty acid incorporation and is synergistic with anti-FASII to inhibit *S. aureus* growth

Since mupirocin leads to down-regulation of phospholipid synthesis genes, it might consequently affect *S. aureus* membrane fatty acid composition. To test this, *S. aureus* Newman was grown in SerFA without and with sublethal mupirocin addition (0.05 μg/ml, *i.e.*, 5-fold below the MIC; (32)). Incorporated eFA was markedly decreased, from 50% in non-treated, to 35% in mupirocin cultures (**Fig. 6A**). (p)ppGpp induction during anti-FASII-induced latency could thus slow or stop eFA incorporation in this transient period.

**Fig. 6.**
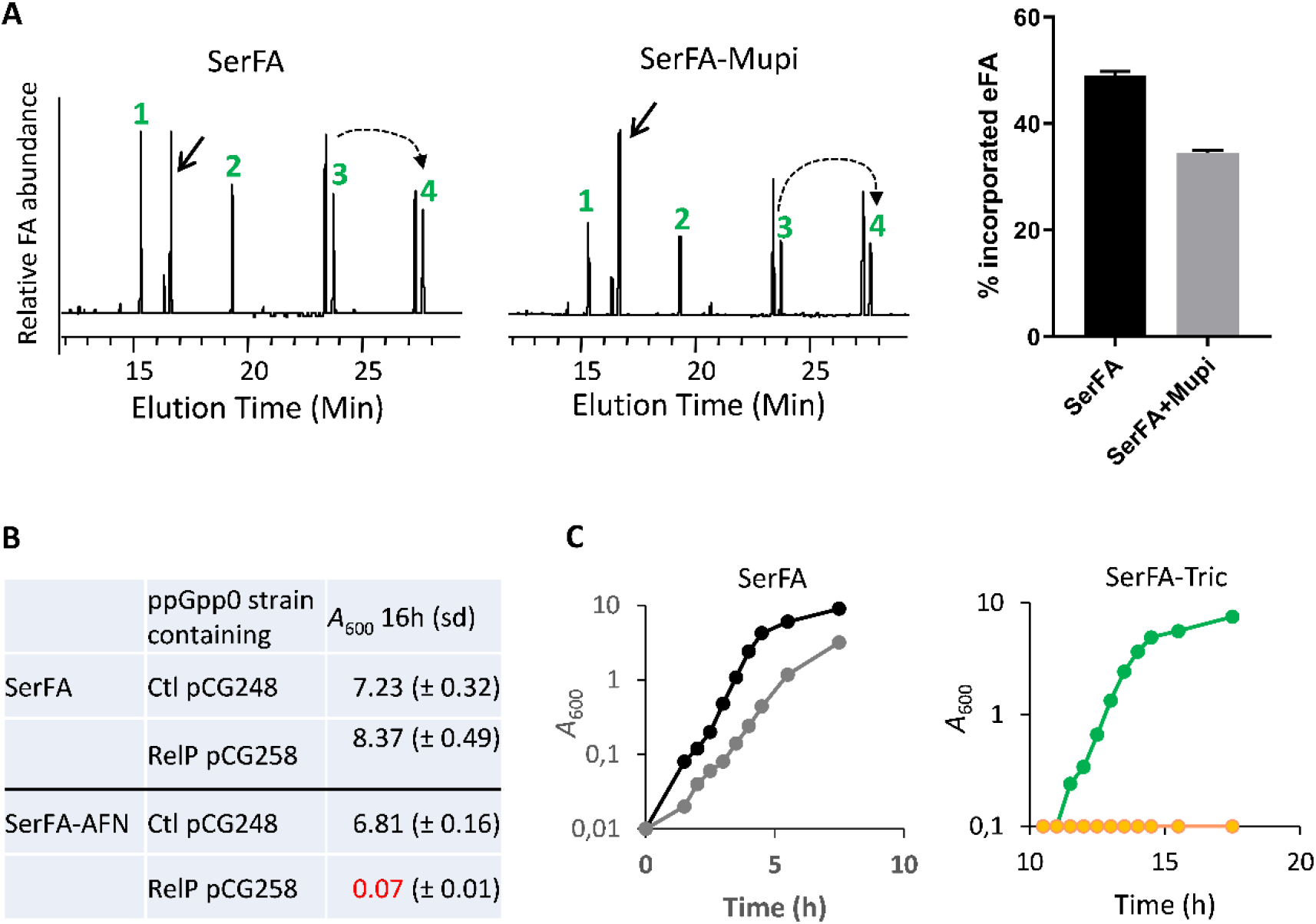
Subinhibitory mupirocin inhibits *S. aureus* eFA incorporation in membrane phospholipids and synergizes with anti-FASII to inhibit adaptation. **A.** Left, Fatty acid profiles of *S. aureus* Newman grown in SerFA and SerFA+mupirocin (mupi). Samples were processed after 3 h growth. Black arrow, position of the main endogenous fatty acid *ai*15:0. eFA are: 1-C14:0; 2-C16:0; 3-C18:1; 4-C20:1 (elongation of C18:1). Dashed arrow, elongation of C18:1 (n+2). Right, percent eFA of Newman grown in SerFA without and with Mupi, derived from integration of two fatty acid profiles from two independent experiments. **B.** Control plasmid pCG248 and aTc-inducible *relP*-expressing plasmid pCG258 (26) were established in the ppGpp0 (null) strain. Strains were grown in SerFA and SerFA-AFN for 16h in the absence of inducer. Anti-FASII adaptation is inhibited in the (p)ppGpp-expressing strain. **C.** Left, *S. aureus* Newman was grown in SerFA (black) and SerFA+mupi (grey), or right, in SerFA-Tric (green) and SerFA-Tric+mupi (orange). Growth was monitored by *A*_600_. Growth curves at right are shown starting at 10 h. Mupi was used at 0.05 μg/ml. Results are representative of three independent experiments.

The above findings led us to hypothesize that FASII inhibitors could be synergistic with a stringent response inducer that prevents compensatory eFA incorporation by repressing phospholipid synthesis genes *plsX* and *plsC*. We first examined anti-FASII adaptation in a strain expressing (p)ppGpp (*via relP*-expressing plasmid pCG258 in a ppGpp0 strain; (26), **Table S2**). While the ppGpp0 control strain (carrying the empty vector pCG248) adapted to anti-FASII after overnight growth, basal RelP expression was sufficient to inhibit anti-FASII adaptation (**Fig. 6B**). This result shows that (p)ppGpp accumulation synergizes with anti-FASII to block *S. aureus* growth. Likewise, addition of a subinhibitory concentration of mupirocin (0.05 μg/ml) and triclosan (0.5 μg/ml) to *S. aureus* SerFA cultures resulted in extended latency, whereas neither mupirocin nor the anti-FASII separately blocked bacterial growth (**Fig. 6C**). Similar results were obtained using anti-FASII AFN-1252 (7) and MRSA strain USA300 FPR3757 (**Table S5**). The observed synergistic effect between two flawed antibiotics may offer an effective strategy for development of last-resort treatments against *S. aureus* infection.

## Discussion

This study reveals the nature of cross-control between *S. aureus* responses to FASII inhibition and to stringent conditions. GTP is depleted in both these conditions, which may explain why the same targets are affected. Our results further show that (p)ppGpp induction lengthens the latency phase preceding adaptation to FASII inhibition*. accBC* transcription is repressed upon stringent response induction, which sets off a chain of events leading to transient repression of phospholipid synthesis genes *plsX* and *plsC*. These events correlate with limited eFA incorporation and extended latency. During *S. aureus* adaptation outgrowth, the initial effects of anti-FASII are reversed, allowing eFA incorporation and adaptation to FASII antibiotics. These results suggest a model (**Fig. 7**) in which (p)ppGpp induction and anti-FASII both initially trigger GTP depletion, resulting in decreased malonyl-CoA pools. The suggested role for CodY in regulating ACC expression remains to be investigated. These events repress phospholipid enzyme synthesis and contribute to anti-FASII latency prior to adaptation outgrowth. Stringent conditions in host niches may be relevant to *S. aureus* infection (33), and might impact the bacterial response to anti-FASII treatment. While our findings identify a role for (p)ppGpp induction *via* GTP depletion in anti-FASII adaptation in *S. aureus*, they do not rule out other roles for these metabolites, or the involvement of other factors in this process.

**Fig. 7.**
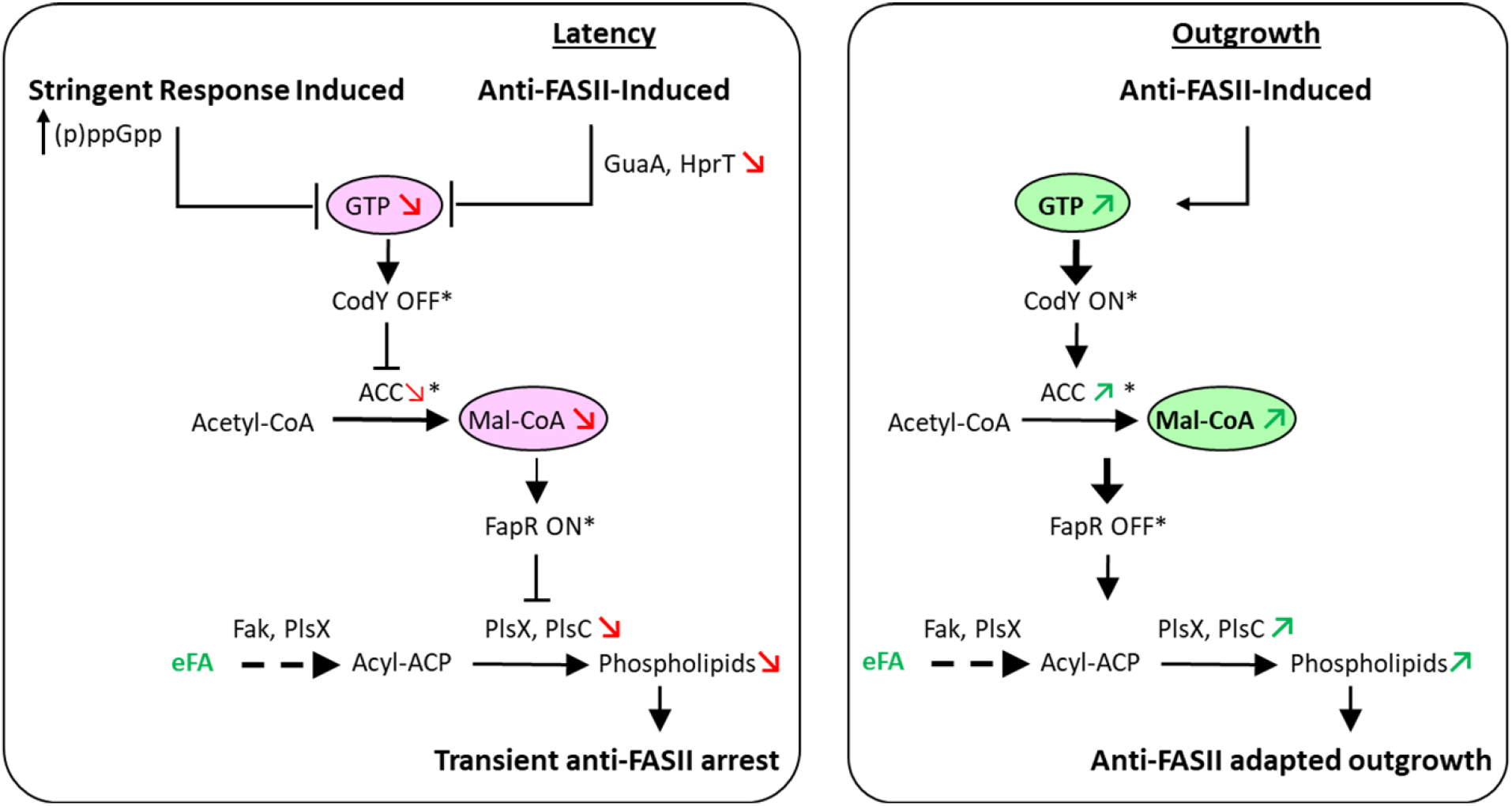
Model for intersecting control of stringent response and FASII inhibition in *S. aureus*. **Left:** Stringent response induction leads to GTP depletion, which in turn modulates gene expression to prepare for starvation (20, 31). We showed that inhibition of FASII also leads to GTP depletion, pointing to an intersecting link between these pathways. Both conditions activate stringent response sensors (P_*ilvD*_-*lacZ*, P_*oppB*_-*lacZ*, and P_*cshA*_-*lacZ*) and lower malonyl-CoA (mal-CoA) pools, such that FapR (for which mal-CoA acts as anti-repressor) exerts repression (13). As a consequence, genes under FapR control, including PlsX and PlsC, remain repressed, blocking phospholipid synthesis. Both stringent response induction and FASII inhibition during the latency phase lead to membrane synthesis arrest. **Right:** Upon FASII adaptation, GTP levels are restored. *accBC* expression is restored to normal, and mal-CoA levels are increased, leading to FapR derepression. In consequence, PlsX and PlsC are both increased, such that phospholipid synthesis resumes. Red and green arrows and circled metabolites correspond to functions analyzed in this study.

A new role for malonyl-CoA in anti-FASII adaptation is uncovered, *via* its increased association with FapR in antibiotic-adapted cultures compared to non-selective cultures. FapR-Trap showed ^~^10-fold greater expression in anti-FASII-adapted cultures than in non-selective cultures, while total malonyl-CoA pools were the same in both conditions (**Fig. 3**). Increased malonyl-CoA interaction with FapR, *i.e.*, FapR derepression, during anti-FASII adaptation is consistent with increased *plsX* and *plsC* expression (**Fig. 5)**. Malonyl-CoA rerouting in anti-FASII may be explained by FabD inactivation in anti-FASII adaptation conditions, e.g., by an intermediate metabolite as suggested in *E. coli* (34). Along this line, a recent study proposed that acyl-ACP accumulation could inhibit FabD (35). Interestingly, acyl-ACP accumulates in a *fabD* mutant during anti-FASII adaptation (4). Alternative possibilities may be considered, such as i- post-translational FabD modification (36) or ii- FabD reversal upon FASII inhibition due to a pile-up of its end-product, malonyl-ACP. We are currently investigating these hypotheses. These findings indicate limits to the reliability of FapR operon-based sensors to estimate malonyl-CoA pools, for which readouts vary according to growth conditions. This may be important to consider in bioengineering applications that rely on FapR operon-like sensors to optimize malonyl-CoA production (37).

Previous studies identified *S. aureus plsC* as comprising a FapR binding site (13). This is disproven here, as FapR failed to bind the published *plsC* consensus site, which lacks a consensus palindromic sequence (**Fig. 5C**). Nevertheless, *plsC* expression is increased in a Δ*fapR* mutant, indicating indirect FapR control of expression.

The need for antimicrobial alternatives is urgent and, besides the discovery of new molecules or targets, the development of efficient combinations based on existing but individually ineffective drugs remains to be explored. Our clarification of the link between stringent response and anti-FASII adaptation opens perspectives for combinatorial antibiotic strategies, using FASII inhibitor and subinhibitory concentrations of stringent response inducers that delay or prevent anti-FASII adaptation of multidrug resistant pathogens like *S. aureus*. Mupirocin, which is usually used topically, was recently proposed as a potentially active systemic antibiotic when presented in liposomes (38). The proof of concept demonstrated here using anti-FASII antibiotics and mupirocin, suggests a useful bi-therapy approach for reducing *S. aureus* survival during infection.

## Materials and Methods

### Strains and media

Strains are listed in **Table S1**. BHI (Brain Heart Infusion) and LB (Luria-Bertani) media were used respectively for *S. aureus* and *E. coli* growth. *S. aureus* precultures were routinely prepared in BHI medium. Three fatty acids (C14:0; myristic acid, C16:0; palmitic acid and C18:1; oleic acid;, Larodan Fine Chemicals, Sweden) were prepared as 100 mM stocks in DMSO and used at final equimolar concentrations of 0.17 mM each in experiments (referred as eFA). Ser-FA (BHI containing eFA+10% newborn calf serum; Sigma) and SerFA-Tric (SerFA+Triclosan 0.5 μg/ml), or SerFA-AFN (SerFA+AFN-1252 0.5 μg/ml) were the modified media used as indicated. Antibiotics kanamycin (50 μg/ml) and erythromycin (5 μg/ml) were used in *E. coli* and *S. aureus* respectively to select pTCV-*lac*-based reporter fusion plasmids (39). Antibiotic adaptation experiments with plasmid-carrying strains were done in SerFA-Tric containing 2 μg/ml erythromycin; note that the latency period was extended by about 4-6 h in this condition. Mupirocin (Sigma), a functional analogue of isoleucyl-AMP and a stringent response inducer of (p)ppGpp (40) was prepared in DMSO and used at 0.1 μg/ml (32, 41) to validate (p)ppGpp sensors, or at 0.05 μg/ml when used in combination with anti-FASII. Equal volumes of DMSO were added to control samples when mupirocin was used.

### Growth experiments in anti-FASII

Three *S. aureus* strains or their derivatives were used to follow anti-FASII adaptation: Newman, USA300, and HG1-R. The latter strain corresponds to HG001 that was repaired for a *fakB1* defect present in the 8325 lineage, and in a minority of *S. aureus* strains; *fakB1* encodes a fatty acid kinase subunit for saturated fatty acid phosphorylation, which enables the use of eFA during anti-FASII adaptation. The above strains showed comparable responses to conditions tested in this work. In experiments using triclosan as anti-FASII, cells were precultured in BHI and then diluted to *A*_600_ = 0.01 in SerFA-Tric. Growth was followed by *A*_600_ readings as indicated. Nonselective exponential and stationary phase cultures were harvested at *A*_600_= 1-4, and ^~^10, respectively. If AFN-1252 was used as anti-FASII, the procedure was the same except that pre-cultures were done in SerFA. Both triclosan and AFN-1252 specifically inhibit FabI, a FASII enzyme (7, 42). Triclosan causes non-specific membrane damage at higher concentrations (42), and was therefore not used in studies with the HG1-R strain, which showed higher resistance to this drug.

### Construction of *fakB1*-repaired strains

The *fakB1* gene of *S. aureus* HG001, as the entire 8325 lineage, displays a 483-bp deletion removing 56% of the 867-bp functional gene. To repair this deletion, a 1,939-bp DNA containing a functional *fakB1* was amplified by PCR from *S. aureus* JE2 with primers fakB1_fp and fakB1_rp (**Table S6**). This fragment was cloned using Gibson Assembly protocol in the thermosensitive vector pG1 (43) amplified with the primers pG1_GibBam and pG1_GibEco (**Table S6**). The resulting vector pG1Ω*fakB1* was introduced by electroporation in *S. aureus* HG001, HG001Δ*fapR*, and HG001ppGpp0, to generate corresponding *fakB1*-repaired strains (**Table S1**). Electroporation of *S. aureus* strains and allelic exchange were performed as described (44). The expected *fakB1* reparation in these strains was confirmed by PCR and sequence analysis.

### Reporter fusions

Promoter regions of *ilvD*, *oppB*, *cshA*, as well as *fapR*, *plsC* and *accBC*, were cloned in pTCV-*lac* or pAW8 plasmids (**Table S2**) using the appropriate primers (**Table S6**). PCR-amplified DNA fragments and plasmid were treated with restriction enzymes EcoRI and BamH1 and ligated products were transformed into DH5α, Top10, or IM08B. The obtained constructs were confirmed by DNA sequencing. Plasmids obtained from IM08B were used directly to transform the *S. aureus* Newman strain; clones obtained in DH5α or Top10 were first established in RN4220. A standard electroporation protocol was used to transform DNA in *S. aureus* (45).

### (p)ppGpp sensors

*ilvD* (NWMN_1960) and *oppB* (NWMN_0856) are up-regulated, and *cshA* (NWMN_1985) genes is down-regulated upon stringent response induction (24, 40, 46). The corresponding stringent response sensors P_*ilvD*_-*lacZ*, P_*oppB*_-*lacZ*, and P_*cshA*_-*lacZ* (**Table S2**) were tested in medium containing 0.1 μg/ml mupirocin (40), and validated as *bone fide* (p)ppGpp sensors (Fig. 1A).

### FapR activity sensor

To estimate malonyl-CoA pools bound to FapR, we designed a transcriptional fusion with promoter and operator sequences containing a consensus FapR binding site; FapR-Trap (**Fig. S2, Table S2**). The construction was based on similar studies done previously to estimate malonyl-CoA pools (37, 47).

### β-galactosidase (β-gal) assay

Fresh cultures were prepared at *A*_600_ = 0.1 from overnight BHI cultures and β-gal activities were measured at the indicated *A*_600_ or time of sampling. When mupirocin was used, cultures were treated or not at *A*_600_ = 0.1 after growth from an initial *A*_600_ = 0.01 and processed 1 h later. All samples of a set were stored at −20°C prior to measurements, which were performed for all samples of a same set. β-gal activities were measured as described (48), except that samples derived from SerFA-containing medium were incubated with lysostaphin (0.1 mg/ml) for 30 min at room temperature prior to processing with β-Glo™ reagents (Promega). β-gal values (mean ± standard deviation) were determined from three independently performed experiments.

### Malonyl-CoA measurement by ELISA assay

Bacterial cultures were prepared as above, and samples were processed at the indicated *A*_600_/time interval according to our test conditions. For each sample, the equivalent of *A*_600_ = 30 was centrifuged at 8,000 rpm at room temperature for 5 min. Pelleted cells were immediately frozen in liquid nitrogen and transferred to −80°C overnight. Ice cold PBS (Phosphate Buffer Saline) was used to resuspend cells at 4°C, which were then sonicated in FastPrep (MP Biomedicals). Supernatants were collected by centrifuging the cell slurry at 13,000 rpm at 4°C for 5 min, and stored at −80°C until use. ELISA tests for total malonyl-CoA measurements were performed as per manufacturer’s instructions (CUSABIO). Malonyl-CoA standards were run along with test samples. Each experiment was performed on three independent cultures. Mean values ± standard deviation are presented.

For malonyl-CoA measurements under stringent response conditions (Fig. 1C, left panel), Newman was first grown in BHI to *A*_600_ of 0.5 from an initial inoculum of 0.01. Cultures were treated or not with 0.1 μg/ml mupirocin for 30 minutes (*A*_600_ = ^~^1 for both samples). ELISA assays were performed as above.

### Purification of *S. aureus* FapR

The *S. aureus fapR* gene was amplified by PCR with FapRORFfp and FapRORFrp primers (**Table S6**) and cloned into pET-21b to produce a recombinant FapR carrying an N-terminal His tag and TEV site expressed in *E. coli* BL21/pDIA17 (13, 49). Bacterial cultures were grown at 37°C in LB containing ampicillin (100 μg/ml) and chloramphenicol (10 μg/ml) until *A*_600_ = 0.6; expression was then performed following addition of IPTG (0.5 mM) at 20°C for 17 hours. Bacteria were harvested by centrifugation (5g wet weight), washed twice in PBS, and resuspended in 30 ml buffer A (50 mM Tris-HCl pH 7,5, 300 mM NaCl, 1 mM DTT), Benzonase^®^ Nuclease (Sigma-Aldrich) and a protease inhibitor cocktail (Roche). Bacteria were lysed by passage through a CF Cell-Disrupter (Constant Systems Ltd) at 4°C. The lysed culture was centrifuged at 46,000 x g for 1 hour and the supernatant was loaded onto a 1-ml Protino^®^ Ni-NTA column (Macherey-Nagel). The protein was eluted with buffer A + 300 mM imidazole and protein-containing fractions were pooled and dialyzed overnight in buffer A with TEV protease (1/10 w/w ratio) at 4°C (produced by the Pasteur Institute Production and Purification of Recombinant Proteins Technological Platform). The His-Tag free protein was loaded onto a 1-ml Ni-NTA column and collected. FapR was further purified using a HiLoad 16/60 Superdex 75 prep grade column (GE Healthcare) equilibrated with 20 mM Tris pH 7.5, 50 mM NaCl. The purified protein was concentrated and stored at −80°C.

### DNase I footprinting

P_*fapRplsX*_ and P_*plsC*_ promoter probes were amplified by PCR from pJJ013 (P_*fapR plsX*_-*lacZ*) and pJJ019 (P_*plsC*_-*lacZ*) (**Table S2**), respectively, with specific promoter primers (P*fapRplsX*_Fw; P*plsC*_fd) and vector primer (pTCV-*lac_*rev).

The 5′ end of pTCV-*lac_*Rev was labeled with (γ-^32^P)ATP, with T4 polynucleotide kinase. Before the DNA binding reaction, purified FapR was dialyzed in 100 mM Na2HPO4/NaH2PO4 (pH 8), 250 mM NaCl, 10 mM MgCl2, DTT 5 mM, and 50% glycerol. DNase I footprinting reactions were performed as described (50). Briefly, 0 to 12 nmol FapR was mixed with 0.2 pmol of DNA and incubated with DNase 1 at room temperature (∼24°C) for 1 min. Samples were analyzed by electrophoresis on a 6% polyacrylamide gel containing 7 M urea. Maxam and Gilbert sequencing ladders (G+A) were loaded on the same gel.

### Determination of *S. aureus* fatty acid profiles

Fatty acid profiles were done as described (4). Newman pre-cultures prepared from two independent colonies were diluted to *A*_600_ = 0.1 in SerFA and grown 3 h with and without mupirocin (0.05 μg/ml). *A*_600_ of SerFA samples were ^~^2.5 and treated samples ^~^1.0. Percentage of eFA are shown (mean ± standard deviation).

### GTP determinations

All extraction steps were performed on ice. Cellular pellets were deproteinized with an equal volume of 6% perchloric acid (PCA), vortex-mixed for 20 s, ice-bathed for 10 min, and vortex-mixed again for 20 s. Acid cell extracts were centrifuged at 13,000 rpm for 10 min at 4°C. The resulting supernatants were supplemented with an equal volume of bi-distilled water, vortex-mixed for 60 s, and neutralized by addition of 2 M Na_2_CO_3_. Extracts were injected onto a C18 Supelco 5 μm (250 × 4.6 mm) column (Sigma) at 45°C. The mobile phase was delivered at a flow-rate of 1 ml/min using the following stepwise gradient elution program: A–B (60:40) at 0 min→(40:60) at 30 min→(40:60) at 60 min. Buffer A contained 10 mM tetrabutylammonium hydroxide, 10 mM KH2PO4 and 0.25% MeOH, and was adjusted to pH 6.9 with 1 M HCl. Buffer B consisted of 5.6 mM tetrabutylammonium hydroxide, 50 mM KH2PO4 and 30% MeOH, and was neutralized to pH 7.0 with 1 M NaOH. Detection was done with a diode array detector (PDA). The LC Solution workstation chromatography manager was used to pilot the HPLC instrument and to process the data. Products were monitored spectrophotometrically at 254 nm, and quantified by integration of the peak absorbance area, employing a calibration curve established with various known nucleosides. Finally, a correction coefficient was applied to correct raw data for minor differences in the total number of cells (by A_600_ measurements) determined in each culture condition.

### Statistical analyses

Graphs and statistical analyses were prepared using GraphPad Prism Software. Mean and standard deviation are presented for sensor fusions, ELISA readouts, fatty acid profile comparisons, and GTP measurements. Statistical significance was determined by unpaired, non-parametric Mann Whitney tests, as recommended for small sample sizes, here biological triplicates, and by a non-parametric, unpaired Kruskal Wallis test for three-way comparisons.

## Acknowledgements

We acknowledge the valuable comments of anonymous reviewers. We are thankful to Jong-In Hong, Seoul National University, Korea, for the generous gift of PyDPA used in initial studies. C. Thomas and members of the Institut Pasteur Production and Purification of Recombinant Proteins Platform are gratefully acknowledged for providing purified FapR. Our colleagues P. Bouloc (I2BC, France), C. Morvan (U. Paris-Descartes, France), and MicrobAdapt team members E. Borezée-Durant, D. Lechardeur, G. Kénanian, R. Boudjemaa, and P. Gaudu gave valuable advice. G. Dubey and N. Descoeudres provided kind support. We acknowledge financial support from DIM-MalInf (AP fellowship), French Agence Nationale de la Recherche (StaphEscape project ANR-13001038) and Fondation pour la Recherche Medicale (DBF20161136769). This work is dedicated to our friend and colleague G. Lamberet, who passed away Dec 23, 2019.

## Author Contributions

Physiology, molecular biology, (p)ppGpp, GTP, and malonyl-CoA estimation: AP, MG, LD, JD, DH, PTC. Fatty Acid analyses: GL, KG and AG. Data analyses: AP, JAM, PTC, and AG. Statistical analysis: DH. Experimental design and project conception: AP, PTC, AG. AG directed the project and AP, KG and AG wrote the manuscript.

## Supplementary information titles and legends

**Supplementary Fig. S1. *S. aureus* bypasses FASII inhibition by exogenous fatty acid (eFA) incorporation in membrane phospholipids. A.** Model for anti-FASII adaptation. FASII and FASII bypass are schematized as characterized; functions whose expression is controlled by FapR repressor are underlined (1–4). Malonyl-CoA reverses FapR repression (5). eFA phosphorylation by Fak (fatty acid kinase) (6) provides an intermediate that may either be incorporated in position 1 of the glycerol-3-phosphate backbone *via* PlsY, or act as a substrate for PlsX to then be incorporated in position 2 via PlsC. In the absence of serum, FabD (malonyl-CoA:ACP transacylase) mutations promote anti-FASII adaptation (2). In contrast, serum favors FASII antibiotic adaptation without FASII mutations (7). **B.**Example of fatty acid profiles of *S. aureus* Newman. Left, BHI grown cells; right, cells grown overnight in SerFA-Tric. Cultures started at *A*_600_ = 0.01 were harvested at *A*_600_ = 1. Arrow indicates *anteiso* 15 (*ai*15), the major fatty acid synthesized by *S. aureus*. eFA: 1, C14:0; 2, C16:0; 3, 18:1. Profiles are representative of three independent experiments. FA, fatty acids; FA-ACP, fatty acyl-ACP; FA-PO_4_, acyl-phosphate; grey, inhibited pathway. FabD^*^, mutated or inhibited enzyme.

**Supplementary Fig. S2. Responses of P**_*ilvD*_-*lacZ***, P**_*cshA*_-*lacZ***, and P**_*accBC*_-*lacZ* **to mupirocin depend on the presence of (p)ppGpp.** HG1-R is an HG001 derivative repaired for a *fakB1* defect common to the 8325 lineage (1). ppGpp0 is the HGR-1 strain devoid of the three synthase genes *rsh*, *relP*, and *relQ* (2). The indicated strains were plated (1 ml of *A*_600_ = 0.1) on BHI medium containing 100 μg/ml X-gal and 5 μg/ml erythromycin and allowed to dry. Mupirocin (75, 37.5, 18.3, and 9.1 ng in rows starting from upper left) was deposited in 3 μl drops. **A**. Expression of P_*ilvD*_-*lacZ* is induced by mupirocin (Fig. 1A), seen as a blue ring in HG1-R, which is absent in the ppGpp0 strain. **B**. P_*cshA*_-*lacZ* is repressed by mupirocin (Fig. 1A), seen as a non-blue growth ring, which is quasi-absent in the ppGpp0 strain. **C.**The P_*accBC*_-*lacZ* sensor behaves like P_*cshA*_-*lacZ*, indicating that production of malonyl-CoA by ACC is repressed by the stringent response. Plates were photographed after 24 h at 37°C and 24 h at 4°C. Dark zones indicate growth inhibition by mupirocin. The ppGpp0 strain is more sensitive to mupirocin than the isogenic parent. Experiments were performed three times giving comparable results.

**Supplementary Fig. S3. FapR-Trap, a malonyl-CoA sensor based on FapR operon** *lacZ* **fusion.**

**A.** Schematic design of FapR-Trap (pJJ004, Table S1). Malonyl-CoA (red diamond) binds FapR (pacman) leading to its release from the FapR binding site (bar code) and expression of *lacZ* (blue) to produce β-galactosidase. The 17 bp FapR consensus binding site used in the construction is shown (based on (1)); converging arrows indicate the 8 bp inverted repeat. **B.** Validation of the FapR-Trap as sensor. β-gal assays were performed with RN4220 and its Δ*fapR* derivative RN4220_Δ*fapR*_ (1) carrying FapR-Trap after 3 h growth in SerFA (left). FapR-Trap expression was also compared in exponential (Exp) and stationary phase (Stat) of the Newman strain (right). Data presented are mean ± standard deviation from triplicate independent experiments. *, p ≤ 0.05 using Mann Whitney.

**Table S1. Strains used in this study.**

**Table S2. Plasmids and constructions.**

**Table S3. Total and proportion of FapR-bound malonyl-CoA depends on growth condition.**

**Table S4. Responses of FapR regulon genes and known stringent response induced genes to mupirocin.**

**Table S5. Subinhibitory mupirocin treatment synergizes with AFN-1252 to inhibits MRSA USA300 growth.**

**Table S6. Primers.**

