## Supplementary Figures and Tables for "(p)ppGpp/GTP and malonyl-CoA modulate *Staphylococcus aureus* adaptation to FASII antibiotics and provide a basis for synergistic bi-therapy"

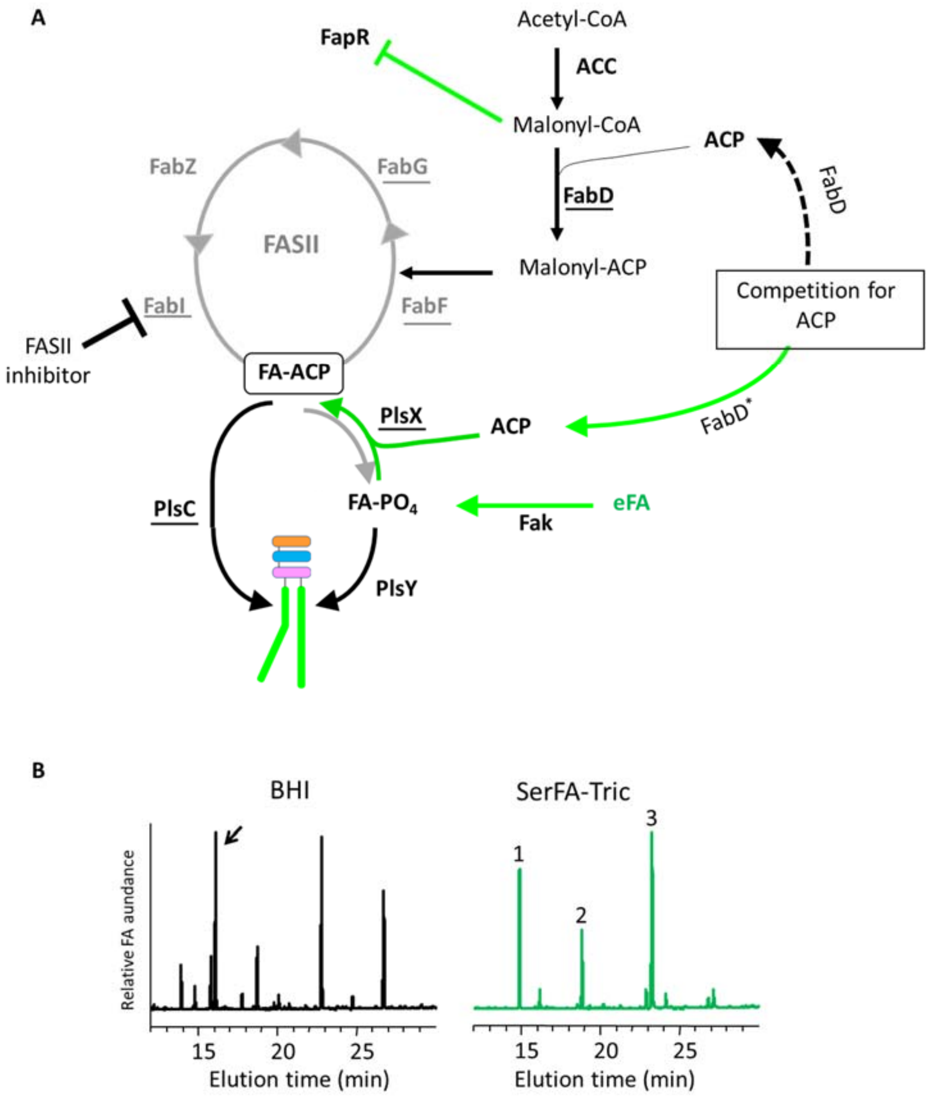


**Supplementary Fig. S1.** ***S. aureus*** **bypasses FASII inhibition by exogenous fatty acid (eFA) incorporation in membrane phospholipids. A.** Model for anti-FASII adaptation. FASII and FASII bypass are schematized as characterized; functions whose expression is controlled by FapR repressor are underlined (1-4). Malonyl-CoA reverses FapR repression (5). eFA phosphorylation by Fak (fatty acid kinase) (6) provides an intermediate that may either be incorporated in position 1 of the glycerol-3-phosphate backbone *via* PlsY, or act as a substrate for PlsX to then be incorporated in position 2 via PlsC. In the absence of serum, FabD (malonyl-CoA:ACP transacylase) mutations promote anti-FASII adaptation (2). In contrast, serum favors FASII antibiotic adaptation without FASII mutations (7). **B.** Example of fatty acid profiles of *S. aureus* Newman. Left, BHI grown cells; right, cells grown overnight in SerFA-Tric. Cultures started at $A_{600}$ = 0.01 were harvested at $A_{600}$ = 1. Arrow indicates *anteiso* 15 (*ai*15), the major fatty acid synthesized by *S. aureus*. eFA: 1, C14:0; 2, C16:0; 3, 18:1. Profiles are representative of three independent experiments. FA, fatty acids; FA-ACP, fatty acyl-ACP; FA-$PO_4$, acyl-phosphate; grey, inhibited pathway. FabD*, mutated or inhibited enzyme.

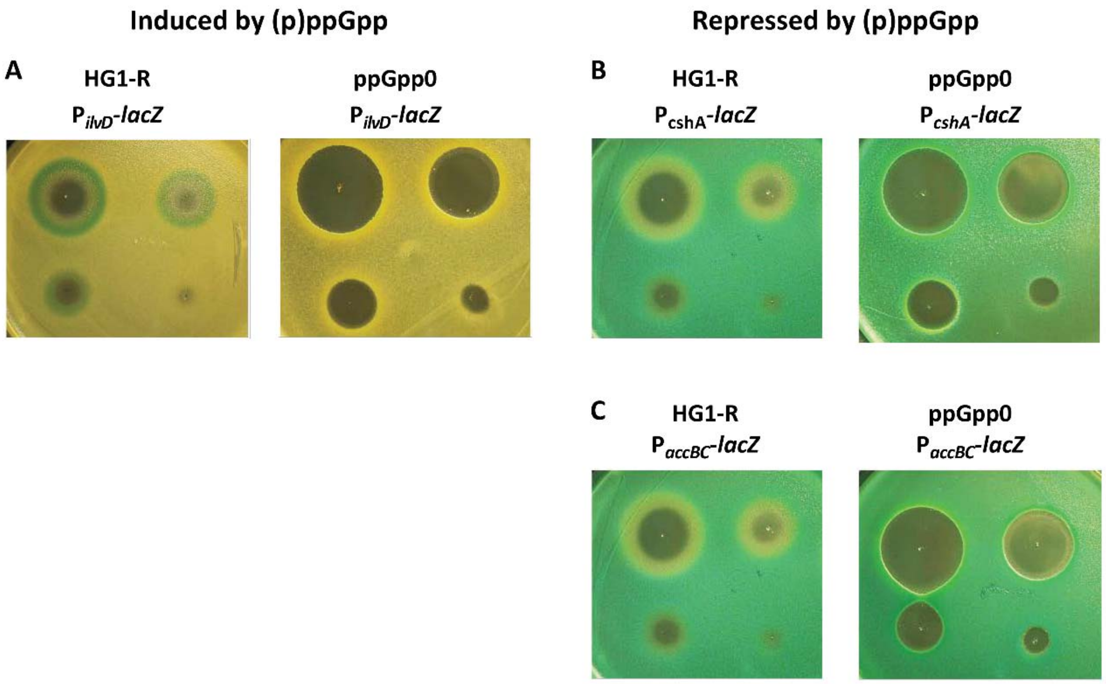


**Supplementary Fig. S2. Responses of $P_{ilvD}$-*lacZ*, $P_{cshA}$-*lacZ*, and $P_{accBC}$-*lacZ* to mupirocin depend on the presence of (p)ppGpp.** HG1-R is an HG001 derivative repaired for a *fakB1* defect common to the 8325 lineage (1). ppGpp0 is the HGR-1 strain devoid of the three synthase genes *rsh*, *relP*, and *relQ* (2). The indicated strains were plated (1 ml of $A_{600}$ = 0.1) on BHI medium containing 100 µg/ml X-gal and 5 µg/ml erythromycin and allowed to dry. Mupirocin (75, 37.5, 18.3, and 9.1 ng in rows starting from upper left) was deposited in 3 µl drops. **A**. Expression of $P_{ilvD}$-*lacZ* is induced by mupirocin (Fig. 1A), seen as a blue ring in HG1-R, which is absent in the ppGpp0 strain. **B**. $P_{cshA}$-*lacZ* is repressed by mupirocin (Fig. 1A), seen as a non-blue growth ring, which is quasi-absent in the ppGpp0 strain. **C.** The $P_{accBC}$-*lacZ* sensor behaves like $P_{cshA}$-*lacZ*, indicating that production of malonyl-CoA by ACC is repressed by the stringent response. Plates were photographed after 24 h at 37°C and 24 h at 4°C. Dark zones indicate growth inhibition by mupirocin. The ppGpp0 strain is more sensitive to mupirocin than the isogenic parent. Experiments were performed three times giving comparable results.

1. Kenanian G, Morvan C, Weckel A, Pathania A, Anba-Mondoloni J, Halpern D, Gaillard M, Solgadi A, Dupont L, Henry C, Poyart C, Fouet A, Lamberet G, Gloux K, Gruss A. 2019. Permissive Fatty Acid Incorporation Promotes Staphylococcal Adaptation to FASII Antibiotics in Host Environments. Cell Rep 29:3974-3982 e4.
2. Geiger T, Kastle B, Gratani FL, Goerke C, Wolz C. 2014. Two small (p)ppGpp synthases in *Staphylococcus aureus* mediate tolerance against cell envelope stress conditions. J Bacteriol 196:894-902.

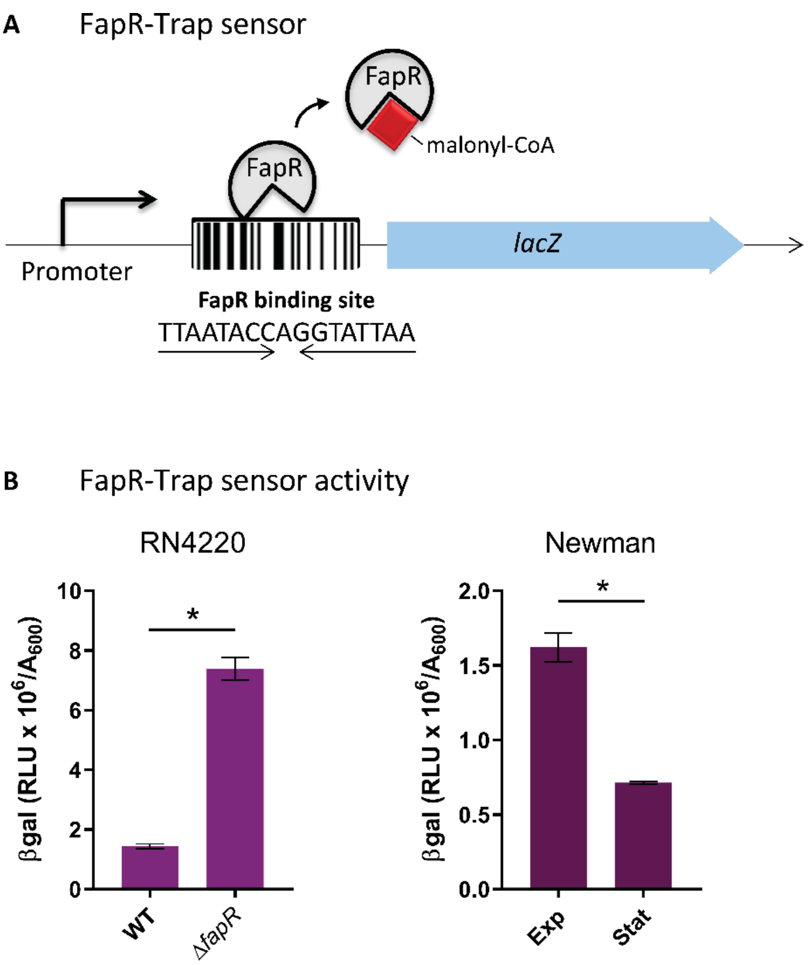


**Supplementary Fig. S3. FapR-Trap, a malonyl-CoA sensor based on FapR operon *lacZ* fusion.** **A.** Schematic design of FapR-Trap (pJJ004, Table S1). Malonyl-CoA (red diamond) binds FapR (pacman) leading to its release from the FapR binding site (bar code) and expression of *lacZ* (blue) to produce β-galactosidase. The 17 bp FapR consensus binding site used in the construction is shown (based on (1)); converging arrows indicate the 8 bp inverted repeat. **B.** Validation of the FapR-Trap as sensor. β-gal assays were performed with RN4220 and its Δ*fapR* derivative RN4220$_{\Delta fapR}$ (1) carrying FapR-Trap after 3 h growth in SerFA (left). FapR-Trap expression was also compared in exponential (Exp) and stationary phase (Stat) of the Newman strain (right). Data presented are mean ± standard deviation from triplicate independent experiments. *, p ≤ 0.05 using Mann Whitney.

1. Albanesi D, Reh G, Guerin ME, Schaeffer F, Debarbouille M, Buschiazzo A, Schujman GE, de Mendoza D, Alzari PM. 2013. Structural basis for feed-forward transcriptional regulation of membrane lipid homeostasis in *Staphylococcus aureus*. PLoS Pathog 9:e1003108.

**Table S1. Strains used in this study.**

| Strain | Phenotypes | Reference |
|---|---|---|
| *S. aureus* | | |
| Newman | Clinical isolate | (1) |
| RN4220 | *S. aureus* cloning recipient ATCC 8325-4 derivative restriction negative | (2) |
| RN4220 Δ*fapR* | RN4220 derivative lacking the FapR repressor of FASII and phospholipid synthesis | (3) |
| USA300 JE2 | USA300_FPR3757 strain devoid of plasmids, derived from methicillin resistant (MRSA) clinical strain | (4) |
| *codY* | USA300_FPR3757 insertional mutant in *codY* (SAUSA300_1148) | (4) |
| HG001 | ATCC 8325 derivative, naturally defective for *fakB1*, a component of the fatty acid kinase. | (1) |
| HG001 Δ*fapR* | Strain lacking the FapR repressor of FASII and phospholipid synthesis | (3, 5, 6) |
| HG1-R | HG001 repaired for defective *fakB1* allele | This study |
| HG1-R Δ*fapR* | HG001 Δ*fapR* repaired for defective *fakB1* allele | This study |
| HG001 ppGpp0 | Triple *rsh*, *relP*, *relQ* mutant that does not produce (p)ppGpp | (7) |
| HG001-R ppGpp0 | HG001 ppGpp0 repaired for defective *fakB1* allele. | This study |
| *E. coli* | | |
| Top10 | F- mcrA Δ(mrr-hsdRMS-mcrBC) φ80lacZΔM15 ΔlacX74 nupG recA1 araD139 Δ(ara-leu)7697 galE15 galK16 rpsL($Str^R$) endA1 $\lambda^-$. | Laboratory collection |
| DH5α | F⁻ *endA1 glnV44 thi-1 recA1 relA1 gyrA96 deoR nupG purB20* φ80d*lacZ*ΔM15 Δ(*lacZYA-argF*)U169, hsdR17($r_K^- m_K^+$), $\lambda^-$ | (8) |
| IM08B | Used for direct cloning in *S. aureus* Newman | (9) |
| BL21 | Used for FapR overexpression | (3) |

1. Herbert S, Ziebandt AK, Ohlsen K, Schafer T, Hecker M, Albrecht D, Novick R, Gotz F. 2010. Repair of global regulators in *Staphylococcus aureus* 8325 and comparative analysis with other clinical isolates. Infect Immun 78:2877-89.
2. Kreiswirth BN, Lofdahl S, Betley MJ, O'Reilly M, Schlievert PM, Bergdoll MS, Novick RP. 1983. The toxic shock syndrome exotoxin structural gene is not detectably transmitted by a prophage. Nature 305:709-12.
3. Albanesi D, Reh G, Guerin ME, Schaeffer F, Debarbouille M, Buschiazzo A, Schujman GE, de Mendoza D, Alzari PM. 2013. Structural basis for feed-forward transcriptional regulation of membrane lipid homeostasis in *Staphylococcus aureus*. PLoS Pathog 9:e1003108.
4. Fey PD, Endres JL, Yajjala VK, Widhelm TJ, Boissy RJ, Bose JL, Bayles KW. 2013. A genetic resource for rapid and comprehensive phenotype screening of nonessential *Staphylococcus aureus* genes. MBio 4:e00537-12.
5. Kenanian G, Morvan C, Weckel A, Pathania A, Anba-Mondoloni J, Halpern D, Gaillard M, Solgadi A, Dupont L, Henry C, Poyart C, Fouet A, Lamberet G, Gloux K, Gruss A. 2019. Permissive Fatty Acid Incorporation Promotes Staphylococcal Adaptation to FASII Antibiotics in Host Environments. Cell Rep 29:3974-3982 e4.
6. Parsons JB, Broussard TC, Bose JL, Rosch JW, Jackson P, Subramanian C, Rock CO. 2014. Identification of a two-component fatty acid kinase responsible for host fatty acid incorporation by *Staphylococcus aureus*. Proc Natl Acad Sci U S A 111:10532-7.
7. Geiger T, Kastle B, Gratani FL, Goerke C, Wolz C. 2014. Two small (p)ppGpp synthases in *Staphylococcus aureus* mediate tolerance against cell envelope stress conditions. J Bacteriol 196:894-902.
8. Sambrook JRDW. 2001. Molecular cloning: a laboratory manual. 3rd ed, Cold Spring Harbor Laboratory, Cold Spring Harbor, NY.
9. Monk IR, Tree JJ, Howden BP, Stinear TP, Foster TJ. 2015. Complete Bypass of Restriction Systems for Major *Staphylococcus aureus* Lineages. MBio 6:e00308-15.

**Table S2. Plasmids and constructions.** [a]

| Plasmid | Description |
|---|---|
| pTCV-*lac* | Promoter expression vector comprising pACYC184 and pAMβ1 as replication origins functional in respectively Gram- negative and positive bacterial hosts (1). |
| pAW8 | *S. aureus* ori-pAMα1–*E. coli* ori-colE1 tetracyline-resistant shuttle vector (2). |
| pG1 | Thermosensitive plasmid used to generate chromosomal gene replacement (3, 4). |
| pG1Ω*fakB1* | pG1 into which a 1,939-bp DNA encoding a functional *fakB1* was cloned by Gibson Assembly. |
| pET-21b | Cloning vector for FapR overexpression in *E. coli* (5). |
| pJJ004 **(FapR-Trap)** | pTCV-*lac* derivative in which *lacZ* transcription is driven by a synthetic promoter designed from the *fabH fabF* (NWMN_0853 and NWMN_0854) operon. An 81 bp hybrid primer-pair (FapRtrapfd and FapRtraprp) was cloned into pTCV-*lac* into EcoRI and BamHI sites. A consensus FapR binding site is present in the promoter (Fig. S2). |
| pJJ005 **($P_{ilvD}$-*lacZ*)** | pTCV-*lac* derivative carrying *lacZ* fused to the promoter region of *ilvD-ilvB* (NWMN_1960 to NWMN_1961). A 515 bp fragment from -500 to +15 with respect to the *ilvD* start codon (genome positions 2168952-2169466) is cloned into pTCV-*lac* EcoRI-BamHI sites. The clone carries an A to G substitution at position-277. This nucleotide change is expected not to alter (p)ppGpp regulation, which occurs *via* CodY (6). |
| pJJ006 **($P_{oppB}$-*lacZ*)** | pTCV-*lac* derivative carrying *lacZ* fused to the promoter region of the *oppB oppC oppD oppF oppA* (NWMN_0856 to NWMN_0860) operon. A 514 bp fragment from -499 to +15 with respect to the *oppB* start codon (genome positions 950296-950809) is cloned into pTCV-*lac* EcoRI-BamHI sites. |
| pJJ008 **($P_{cshA}$-*lacZ*)** | pTCV-*lac* derivative carrying *lacZ* fused to the *cshA* (NWMN_1985) promoter region. A 515 bp fragment from -500 to +15 with respect to the *cshA* start codon (genome positions 2207332-2207846) is cloned into pTCV-*lac* EcoRI-BamHI sites. |
| pJJ013 **($P_{fapR\ plsX}$-*lacZ*)** | pTCV-*lac* derivative carrying *lacZ* fused to the promoter region of the *fapR plsX fabD fabG* (NWMN_1138 to NWMN_1141) operon. A 315 bp fragment from -300 to +15 with respect to the start codon (genome |

| | |
|---|---|
| | positions 1247750-1248064) of the operon is cloned into pTCV-*lac* EcoRI-BamHI sites. |
| pJJ019 (**P*plsC*-*lacZ***) | pTCV-*lac* derivative carrying *lacZ* fused to the *plsC* (NWMN_1620) promoter region. A 315 bp fragment from -300 to +15 with respect to the *plsC* start codon (genome positions 1799913-1800227) is cloned into pTCV-*lac* EcoRI-BamHI sites. |
| pJJ027 (**P*accBC*-*lacZ***) | pTCV-*lac* derivative carrying *lacZ* fused to the promoter region of the *accB accC* (NWMN_1432, NWMN_1431) operon. Hybrid primer-pair of 103 bp (accBCfp and accBCrp), corresponding to genome positions 1604036-1603934**,** was used to clone the promoter region upstream of *accB* into pTCV-*lac* into EcoRI and BamHI sites. |
| pJJ042 (FapR-ORF) | pET21-b derivative carrying *fapR* ORF (NWMN_1138). An N-terminus hexa-histidine tag followed by a TEV cleavage site is fused to FapR. Here, Nhe1 and Sal1 sites are used for cloning. Cloning strategy is as described(5)**.** |
| pJJ043 (P*accBC*-*lacZ*) | A pAW8-modified derivative where the *accBC-lacZ* fusion was amplified from pJJ027 and cloned into EcoR1 and Sma1 sites of pAW8. |

[a] Designations of promoter fusions are in bold (in parentheses). See Table S6 for primer pairs used for clonings.

**Table S3. Total and proportion of FapR-bound malonyl-CoA depends on growth condition.**

| | Relative proportions [a] | | % Malonyl-CoA bound to FapR |
|---|---|---|---|
| | Total Malonyl-CoA pools | FapR-Trap | |
| Non- selective [b] | 100 | 8 | 8% |
| Anti-FASII [c] Latency | 23 | 10 | 43 % |
| Anti-FASII [c] Adapted- Exponential | 93 | 100 | 108 % |

[a] Relative proportions of total and FapR-bound malonyl-CoA from Fig. 3B and 3C are determined based on the highest obtained value for respective measurements (indicated as 100 %). [b] Medium was SerFA. [c] FASII inhibitor triclosan is added to SerFA (SerFA-Tric).

**Table S4. Responses of FapR regulon genes and known stringent response induced genes to mupirocin.**

| | Promoter fusion | Mupirocin/ No addition [a] | Effect |
|---|---|---|---|
| Control | Pctl [b] | 1.1 ± 0.03 | None |
| Stringent response | $P_{ilvD}$-*lacZ* | 2.1 ± 0.42 | Stimulation |
| | $P_{oppB}$-*lacZ* | 1.87 ± 0.69 | Stimulation |
| | $P_{cshA}$-*lacZ* | 0.33 ± 0.06 | Repression |
| FapR regulon | $P_{accBC}$-*lacZ* | 0.25 ± 0.04 | Repression |
| | $P_{fapR\ plsX}$-*lacZ* | 0.25 ± 0.03 | Repression |
| | $P_{plsC}$-*lacZ* | 0.34 ± 0.06 | Repression |

[a] Measurements (standard deviation) were determined on three independent samples in BHI medium containing or not mupirocin 0.1 µg/ml (described in Materials and Methods). Experiments were performed independently from those presented in Fig. 1. [b] Pctl corresponds to the plasmid vector pTCV-lac (1), lacking a *lacZ* promoter.

1. Poyart C, Trieu-Cuot P. 1997. A broad-host-range mobilizable shuttle vector for the construction of transcriptional fusions to beta-galactosidase in gram-positive bacteria. FEMS Microbiol Lett 156:193-8.

**Table. S5. Subinhibitory mupirocin treatment synergizes with AFN-1252 to inhibits MRSA USA300 growth.** [a]

| **Media** | **$A_{600}$ at 16 h** |
|---|---|
| SerFA | 11.4 ±0.3 |
| SerFA+AFN-1252 [b] | 11.1 ±0.7 |
| SerFA + Mupirocin | 6.3 ±1.7 |
| SerFA-AFN-1252+Mupirocin | 0.2 ±0.06 |

[a] Mupirocin was used at 0.06 µg/ml, AFN-1252 at 0.5 µg/ml. The USA300_FRPR3757 strain is MRSA (methicillin resistant *S. aureus*). It was precultured in SerFA medium, and diluted 1:100 in SerFA containing or not mupirocin and AFN-1252. $A_{600}$ optical densities were determined after 16 h aerobic growth at 37°C. Results shown are the average (range) of 2 independent experiments. [b] AFN-1252 is a pipeline antibiotic that, like triclosan, targets FabI (1).

1. Banevicius MA, Kaplan N, Hafkin B, Nicolau DP. 2013. Pharmacokinetics, pharmacodynamics and efficacy of novel FabI inhibitor AFN-1252 against MSSA and MRSA in the murine thigh infection model. J Chemother 25:26-31.

**Table S6. Primers.** [a]

| Primer ID | Sequence of Primer (5'to 3') |
|---|---|
| FapRtrapfd | AATTCTCCTACGACAATATCCT**TTGCAT**TTAAAGTATCTAAAATC**TATGAT**AAA***TTAATACCTGGTATTAA***AAATATTTATTAGAAG<br>*pJJ004 (FapR-Trap)* |
| FapRtraprp | GATCCTTCTAATAAATATTT***TTAATACCAGGTATTAA***TTT**ATCATA**GATTTTAGATACTTTAA**ATGCAA**AGGATATTGTCGTAGGAG<br>*pJJ004 (FapR-Trap)* |
| ilvDfp | TTGCCGGAATTCCCTATATTATGCTTTTCATTCA<br>*pJJ005 ($P_{ilvD}$-lacZ)* |
| ilvDrp | CTAGCGGGATCCTTACATGTCGCTTCGCATAGT<br>*pJJ005 ($P_{ilvD}$-lacZ)* |
| OppBfp | TTGCCGGAATTCTTGAAAAATGGATCATCAGA<br>*pJJ006 ($P_{oppB}$-lacZ)* |
| OppBrp | CAAGCGGGATCCTTAAATATATTTCCCCATCTAA<br>*pJJ006($P_{oppB}$-lacZ)* |
| CshAfp | TTGCCGGAATTCCTTTTACTTATAAAAATGATTTG<br>*pJJ008 ($P_{cshA}$-lacZ)* |
| CshArp | CAAGCGGGATCCTTATTTAAAATTTTGCAAAATAATTC<br>*pJJ008 ($P_{cshA}$-lacZ)* |
| FapRfp | TTGCCGGAATTCCAGCTGAACTTATTCAATCTGG<br>*pJJ013 ($P_{fapR\ plsX}$-lacZ)* |
| FapRrp | CAAGCGGGATCCTTACGTCTCACCCCTCATTTTTTAGT<br>*pJJ013 ($P_{fapR\ plsX}$-lacZ)* |
| PlsCfp | TTGCCCGAATTCAAGTGCACCAATAATTCCAGCA<br>*pJJ019 ($P_{plsC}$-lacZ)* |
| PlsCrp | CAAGCGGGATCCTTAAATCACTGAATACATTGTGCCACC<br>*pJJ019 ($P_{plsC}$-lacZ)* |
| AccBCfp | AATTCGTCGCCAGCAATATGAACATGC**TTGAAT**TGAAGAGTTGTCTCAAG**TAAAAT**AGACGGGTAGATGAAAACAAACTGAAGGAGTCAGTAATAATGAACTTTAAAG<br>*pJJ027 ($P_{accBC}$-lacZ)* |
| AccBCrp | GATCCTTTAAAGTTCATTATTACTGACTCCTTCAGTTTGTTTTCATCTACCCGTCT**ATTTTA**CTTGAGACAACTCTTCA**ATTCAA**GCATGTTCATATTGCTGGCGACG<br>*pJJ027 ($P_{accBC}$-lacZ)* |
| FapRORFfp | AACTAGCTAGCCATCATCATCATCATCACGAAAACCTGTATTTTCAGGGCATGAGGGGTGAGACGTTGAAACTAAAGAAAG pJJ042 (FapR-ORF) |
| FapRORFrp | AACGCGTCGACTTATCCTCGCTTATCATAAAACATTTTAAAATTTCC<br>pJJ042 (FapR-ORF) |
| pAW8AccBCfp | TTGCCGGAATTCGTCGCCAGCAATATGAACATGCTTG<br>pJJ043 (P*accBC-lacZ* for pAW8 insertion) |
| pAW8AccBCrp | TGCGTAAGGAGAAAATACCGCATCAGCCCGGGTTATTTTTGACACCAGACCAACTG |

| | |
|---|---|
| | pJJ043 (P*accBC-lacZ* for pAW8 insertion) |
| fakB1_fp | CCTGCAGGTCGACTCTAGAGGATCCGTTCGACACGCCCGATATCA |
| fakB1_rp | ACAGCTATGACATGATTACGAATTCTACCTGCACCTGTTACGGC |
| pG1_GibBam | GGATCCTCTAGAGTCGACCTGCAGG |
| pG1_GibEco | GAATTCGTAATCATGGTCATAGCTG |
| *PfapRplsX*_Fw | CGACTAAATAATAGCTAAATATTACAG |
| P*plsC*_Fw | CAACTTTAGATGTATTTTCAGACTATC |
| pTCV-*lac*_Rev (vector primer) | CCACAGTAGTTCACCACCTTTTCCC |

[a] Restriction sites and overlap sequences in Gibson clonings are underlined; putative -10 and -35 motifs are in bold; FapR consensus binding site is in italics.
